# Urinary biomarkers and cardiovascular outcomes in the UK Biobank: observational and Mendelian randomization analyses

**DOI:** 10.1101/499483

**Authors:** Daniela Zanetti, Helene Bergman, Stephen Burgess, Themistocles L. Assimes, Vivek Bhalla, Erik Ingelsson

**Affiliations:** Department of Medicine, Division of Cardiovascular Medicine, Stanford University School of Medicine, Stanford, CA; Stanford Cardiovascular Institute, Stanford University, Stanford, CA; Stanford Diabetes Research Center, Stanford University, Stanford, CA; Department of Medical Epidemiology and Biostatistics, Karolinska Institute, Sweden; Department of Public Health and Primary Care, University of Cambridge, Cambridge, UK; Palo Alto VA Health Care System, Palo Alto, CA; Department of Medicine, Division of Nephrology, Stanford University School of Medicine, Stanford, CA

**Keywords:** urinary biomarkers, kidney function, blood pressure, type 2 diabetes, cardiovascular diseases, Mendelian randomization

## Abstract

**Background:** Urinary biomarkers are associated with hypertension and cardiovascular disease (CVD), but the nature of these associations is incompletely understood.

**Methods:** We performed multivariable-adjusted regression models to assess associations of urinary sodium-potassium ratio (UNa/UK), and urinary albumin adjusted for creatinine (UAlb/UCr) with cardiovascular risk factors, CVD and type 2 diabetes (T2D) in 478,311 participants of the UK Biobank. Further, we studied above associations separately in men and women, and assessed the causal relationships of these kidney biomarkers with cardiovascular outcomes using the two-sample Mendelian randomization (MR) approach.

**Results:** In observational analyses, UNa/UK showed significant inverse associations with atrial fibrillation (AF), coronary artery disease (CAD), ischemic stroke, lipid-lowering medication and T2D. In contrast, UAlb/UCr showed significant positive associations with AF, CAD, heart failure, hemorrhagic stroke, lipid-lowering medication and T2D. We found a positive association between UNa/UK and albumin with blood pressure (BP), as well as with adiposity-related measures. Generally, we detected consistent directionality in sex-stratified analyses, with some evidence for sex differences in the associations of urinary biomarkers with T2D and obesity. After correcting for potential horizontal pleiotropy, we found evidence of causal associations of UNa/UK and albumin with systolic BP (beta_SBP_≥2.63; beta_DBP_≥0.85 SD increase in systolic BP per SD change UNa/UK and UAlb/UCr; *P*≤0.038), and of albumin with T2D (odds ratio=1.33 per SD change in albumin, *P*=0.023).

**Conclusion:** Our Mendelian randomization analyses mirror and extend findings from randomized interventional trials which have established sodium intake as a risk factor for hypertension. In addition, we detect a feed-back causal loop between albumin and hypertension, and our finding of a bidirectional causal association between albumin and T2D reflects the well-known nephropathy in T2D.

## Introduction

Cardiovascular disease (CVD) is a leading cause of mortality worldwide. In 2015, it accounted for 17.9 million deaths, or almost one third of all deaths globally.^1^ New strategies to prevent CVD are highly sought after from both a humanitarian and economical perspective. The identification of causal risk factors associated with CVD is expected to provide important insights on how to develop new strategies.

Over the past decades, several kidney-related biomarkers have been proposed to be associated with CVD,^2,3^ but their potential causal role in disease processes is incompletely understood. We used urinary biomarkers, as proxies for kidney function, to try to shed light on this relationship and to pinpoint causal risk biomarkers involved in this association.

The association of high sodium and low potassium intake with elevated blood pressure is supported by a large body of evidence that includes randomized clinical trials,^4,5^ as well as observational cohort studies.^6,7^ The Trials of Hypertension Prevention (TOPH), conducted in 1987-1990 and 1990-1995, respectively, showed that reducing dietary sodium in individuals with prehypertension decreased the risk of cardiovascular events and overall mortality up to 20 years after the original trial.^8–10^ For CVD, two previous meta-analyses of prospective cohort studies detected significant positive relationships between sodium intake and stroke, as well as CVD outcomes.^11,12^ Similarly, in observational epidemiologic studies, high albuminuria is associated with risk for cardiovascular events in individuals with or without diabetes mellitus.^13,14^ One recent Mendelian randomization (MR) study^15^ supported the existence of a bidirectional causal association between albuminuria and blood pressure, implying that albuminuria could increase risk of cardiovascular disease through blood pressure. There is a lack of prior studies comprehensively examining several urinary biomarkers reflecting different aspects of kidney function and their associations with blood pressure, type 2 diabetes (T2D) and CVD in a large study sample from the general population. Furthermore, the causal associations of these biomarkers with cardiometabolic traits are incompletely understood.

The aims of this study were to first determine the associations between urinary biomarkers, specifically urinary sodium, potassium and albumin, with cardiovascular risk factors, T2D and CVD in the UK Biobank by means of observational analyses including sex-stratified analyses; and then to test whether any of these associations are causal using a two-sample MR approach, combining UK Biobank data with publicly available data from relevant genome-wide association studies (GWAS).

## Methods

### Study Population

The UK Biobank is a longitudinal cohort study of >500 000 individuals aged 40 to 69 years initiated in the United Kingdom in 2006–2010.^16^ We used the data collected at the UK Biobank assessment centers at baseline, combined with information on incident events from the hospital and death registries. In our main analysis (N = 478,311), we excluded participants with diagnoses indicating impaired kidney function (N =7,221) and/or CVD at baseline (atrial fibrillation [AF], coronary artery disease [CAD], heart failure [HF], hemorrhagic stroke [HS], or ischemic stroke [IS]) (N = 17,087; see below). We defined impaired kidney function as International Classification of Diseases (ICD) edition 9 codes 581-589, 591, 2503 and V420; edition 10 codes N00, N01, N03-N08, N10-N19, N25-29, E10.2, E11.2, E14.4, and Z99.2; and surgical codes for kidney (codes M01-M06, M08). In our sensitivity analysis (N = 390,893), we additionally excluded participants using diuretics, angiotensin-converting-enzyme (ACE) inhibitors, angiotensin II receptor blockers (ARBs), or calcium channel blockers (CCB) for any reason, and medications that are combination drugs including one or more of these categories (N = 87,418), as these medications may impact kidney function and therefore urinary excretion of sodium or potassium, or may impact the effect of dietary intake on CVD. Details of these measurements can be found in the UK Biobank Data Showcase (http://biobank.ctsu.ox.ac.uk/crystal/).

### Definition of Exposure and Cardiovascular Outcomes for Observational Analyses

The exposures of interest for our main analysis were urinary sodium (field ID 30530) to potassium (field ID 30520) excretion ratio (UNa/UK), and urinary albumin (field ID 30500) to creatinine (field ID 30510) ratio (UAlb/UCr). We analyzed urinary sodium to creatinine ratio (UNa/UCr) and urinary potassium to creatinine ratio (UK/UCr) for secondary analyses. The method of using spot-urine samples to approximate 24-hour excretion is widely used, as a convenient alternative in assessing population mean sodium and potassium intake, especially for surveys with large populations.^17,18^

Cardiovascular outcomes were defined using the inpatient hospital and death registries, including primary and secondary causes to maximize power. AF was defined as ICD-9 code 427.3, ICD-10 code I48, and surgical codes K50.1, K62.2-K62.4. CAD was defined as ICD edition 9 codes 410-411, edition 10 codes I20.0, I21, and I22; and surgical codes for percutaneous transluminal coronary angioplasty and coronary artery bypass graft (codes K40-K46, K49-K50, and K75). HF was defined as ICD-9 code 428 and ICD-10 code I50. Stroke was defined as hemorrhagic (ICD-9: 430-432, ICD-10: I60-I62) or ischemic stroke (ICD-9: 433-434, ICD-10: I63). The hospital registry-based follow-up ended on March 31, 2015 in England; August 31, 2014 in Scotland; and February 28, 2015 in Wales. We censored individuals either on these dates, at the time of event in question, or at the time of death, whichever occurred first. The death registry included all deaths that occurred before January 31, 2016 in England and Wales, and November 30, 2015 in Scotland.

### Definition of Confounders for Observational Analyses

We used data from questionnaires to derive the following potential confounders: sex (ID 31), age (field ID 21003), region of the UK Biobank assessment center (ID 54; recoded to three countries: UK, Scotland and Wales), ethnicity (ID 21000; recoded to four groups: black, Asian, white, mixed), smoking status (ID 20116, recoded to three groups: never, previous, current), alcohol use (ID 100022, weekly alcohol intake in grams), degree of physical activity (ID 894, recoded to two groups: days/week of moderate physical activity <5, days/week of moderate physical activity ≥5), and a Townsend index reflecting socioeconomic status (ID 189). Physical measurements were used to define systolic blood pressure (SBP) (ID 4080, but if missing ID 93), diastolic blood pressure (DBP) (ID 4079, but if missing ID 940), body fat percentage (ID 23099), body mass index (BMI) (ID 21001), and waist-to-hip ratio (WHR) (ID 48/ID 49). Lipid medications (ID 20003; including the following medications: simvastatin, pravastatin, fluvastatin, atorvastatin, rosuvastatin, ezetimibe, nicotinic acid product or fenofibrate) were used as a proxy for hyperlipidemia, as lipid level measurements were not available in UK Biobank at the time of the present study. T2D was defined as having a diagnosis of ICD-9 code 250.10 or 250.12, or ICD-10 code E11 in the in-patient hospital register; diabetes diagnosed by a physician (ID 2443) after 35 years old (ID 2976), or being treated with anti-diabetic medication, but without insulin treatment in the first year (ID 2986).

### Statistical Methods

#### Observational Analysis

After examining distributions of all variables, we applied rank-based inverse normal transformation for the urinary biomarkers, and for all the continuous variables included in the study. Multivariable-adjusted Cox proportional hazards models were performed to determine associations of our exposures with AF, CAD, HF, HS and IS events, separately; during a median follow-up time of 6.1 years. We performed multivariable linear regression models to determine associations of exposures with SBP, DBP, body fat percentage, BMI, and WHR; and multivariable logistic regression models to study associations of urinary biomarkers with lipid medications and T2D. We assessed evidence of nonlinear effects of UNa/UK and UAlb/UCr on different outcomes using spline regression models. We use the DAGitty web tool (http://dagitty.net/dags.html) to systematically construct our multivariable model adjusting for confounders. All association analyses were adjusted for age, sex, region of the UK Biobank assessment center, ethnicity, smoking, alcohol, physical activity, Townsend index, blood pressure [DBP and SBP], obesity [BMI, body fat percentage, WHR], lipid medications, T2D, and medications affecting renal excretion. Further, we performed sex-stratified analyses to study sex differences of these associations. A Bonferroni-corrected threshold of 4.17E-03 (adjusting for 12 comparisons) was used to identify significant associations. Linear/logistic and Cox regressions analyses were conducted with the R software and R package *Survival* (version 3.3.0), respectively.

#### Mendelian Randomization

We performed two-sample MR analyses using data from publicly available consortia, except for blood pressure where we performed a GWAS in UK Biobank (as the publicly available GWAS summary statistics were adjusted for BMI). We used the results of previous GWAS of UNa/UK, UNa/UCr, UK/UCr and UAlb/UCr performed by our group^19^ available at GRASP resource (https://grasp.nhlbi.nih.gov/FullResults.aspx), as instrument variables (IVs). A list of the variants included in the IV is shown in eTable 1. We assessed the causal relationships of the four urinary biomarkers with risk factors for CVD (SBP, DBP, BMI, WHR). We did not study causal associations of UNa/UK, UNa/Cr and UK/Cr with any hard CVD endpoints due to lack of statistical power (eTable 2). We assessed the causal relationships of UAlb/UCr with AF and T2D (power estimates >75%) (eTable 2).

We performed two-sample MR using three separate methods to estimate causal effects: the standard inverse-variance weighted (IVW) regression; as well as two robust regression methods, the weighted median-based method, and Egger regression.^20^ We performed leave-one-out sensitivity analyses to identify if a single SNP was driving an association. In addition, we performed bidirectional MR and multivariable MR for significant causal outcomes.

We performed the two-sample MR analyses,^20,21^ as well as the bidirectional MR and the multivariable MR with the R package, *TwoSampleMR*. In addition, we used the MR-PRESSO software^22^ to minimize the risk of horizontal pleiotropy affecting our results. Details of the GWAS summary statistics used to performed MR analyses and variance explained by our instruments can be found in eTable 2.

We estimated statistical power for each of the MR analyses using variance explained from our exposures and effect size from observational analyses and an alpha threshold of 0.05. In addition, we calculated the statistical power using a fixed effect of 1.15 for binary trait and 0.15 for continuous traits. Power for MR analyses was estimated with the online tool at https://sb452.shinyapps.io/power/.

## Results

Baseline characteristics of UK Biobank participants are shown in Table 1. In the main analysis, the mean age at baseline was 56.3 years (SD, 8.1 years) and 56% of participants were females. During a median follow-up time of 6.1 years, 22,212 incident CVD cases occurred in participants free from the disease at baseline (9,196 AF; 7,375 CAD; 2,775 HF; 978 HS and 1,888 IS events) (Table 2).

**Table 1.** Baseline characteristics of UK Biobank participants for main (N = 478,311) and sensitivity (N = 390,893) analyses.

|  | Main cohort* | Cohort for sensitivity analyses† |
| --- | --- | --- |
| Variables | N (%) or Mean (SD) | N (%) or Mean (SD) |
| Sex: female | 265,777 (56) | 222,152 (57) |
| Age | 56.3 (8.1) | 55.4 (8.1) |
| Ethnicity: Black | 7,773 (2) | 5,632 (2) |
| Ethnicity: Asian | 10,754 (2) | 8,566 (2) |
| Ethnicity: White | 449,961 (94) | 368,525 (94) |
| Ethnicity: Mixed | 7,207 (2) | 6,013 (2) |
| Smoking: Previous | 161,803 (34) | 126,636 (33) |
| Smoking: Current | 50,067 (11) | 42,434 (11) |
| Weekly alcohol intake, g | 128.6 (159.1) | 127.7 (156.2) |
| Days/week of moderate physical activity <5 | 274,641 (58) | 224,396 (57) |
| SBP, mmHg | 140 (20) | 138 (19) |
| DBP, mmHg | 82 (11) | 82 (11) |
| Body Fat, % | 31.5 (8.6) | 31.0 (8.5) |
| BMI, kg/m<sup>2</sup> | 27.4 (4.8) | 26.8 (4.5) |
| WHR | 0.87 (0.09) | 0.86 (0.09) |
| Lipid medications | 67,898 (14) | 28,978 (7) |
| T2D | 19,876 (4) | 7,758 (2) |
| Medications affecting renal excretion | 87,417 (18) |  |
| Urine Sodium, millimole/L | 77.47 (44.51) | 76.63 (44.40) |
| Urine Potassium, millimole/L | 63.01 (33.88) | 63.10 (34.15) |
| Urine Albumin, mg/L | 13.11 (57.38) | 11.27(41.85) |
| Urine Creatinine, millimole/L | 8.83 (5.80) | 8.81 (5.78) |
| Urine Sodium/Potassium ratio | 1.44 (0.91) | 1.41 (0.87) |
| Urine Sodium/Creatinine ratio | 10.76 (6.98) | 10.53 (6.60) |
| Urine Potassium/Creatinine ratio | 8.41 (4.30) | 8.45 (3.98) |
| Urine Albumin/Creatinine ratio | 16.59 (60.40) | 14.84 (44.84) |

\* Main cohort (N= 478,311) excluded participants with diagnosis indicating decreased kidney function (N=7,221) and/or cardiovascular disease at baseline (N=17,087).
† Cohort for sensitivity analysis (N= 390,893) further excluded participants with specified medications affecting renal excretion (N=87,418).
Abbreviations: SD, standard deviation; T2D, type 2 diabetes; SBP, systolic blood pressure; DBP, diastolic blood pressure; BMI, body mass index; WHR, waist-to-hip ratio.

**Table 2.**
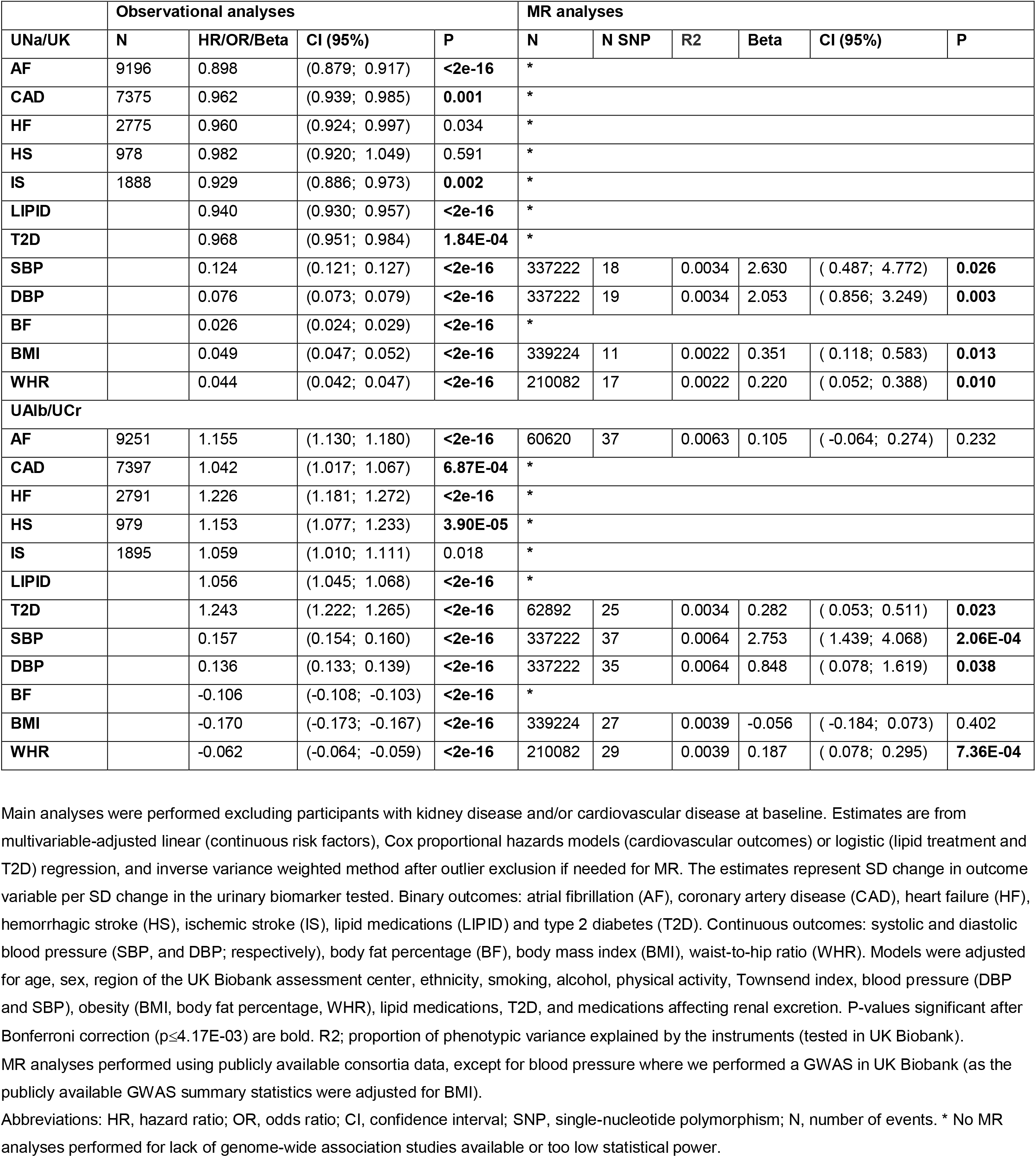
Observational and Mendelian randomization (MR) analyses of associations of urinary sodium-potassium ratio (UNa/UK) and urinary albumin-creatinine ratio (UAlb/UCr) with cardiovascular outcomes.

|  | Observational analyses |  |  |  | MR analyses |  |  |  |  |  |
| --- | --- | --- | --- | --- | --- | --- | --- | --- | --- | --- |
| UNa/UK | N | HR/OR/Beta | CI (95%) | P | N | N SNP | R2 | Beta | CI (95%) | P |
| AF | 9196 | 0.898 | (0.879; 0.917) | **<2e-16** | * |  |  |  |  |  |
| CAD | 7375 | 0.962 | (0.939; 0.985) | **0.001** | * |  |  |  |  |  |
| HF | 2775 | 0.960 | (0.924; 0.997) | 0.034 | * |  |  |  |  |  |
| HS | 978 | 0.982 | (0.920; 1.049) | 0.591 | * |  |  |  |  |  |
| IS | 1888 | 0.929 | (0.886; 0.973) | **0.002** | * |  |  |  |  |  |
| LIPID |  | 0.940 | (0.930; 0.957) | **<2e-16** | * |  |  |  |  |  |
| T2D |  | 0.968 | (0.951; 0.984) | **1.84E-04** | * |  |  |  |  |  |
| SBP |  | 0.124 | (0.121; 0.127) | **<2e-16** | 337222 | 18 | 0.0034 | 2.630 | ( 0.487; 4.772) | **0.026** |
| DBP |  | 0.076 | (0.073; 0.079) | **<2e-16** | 337222 | 19 | 0.0034 | 2.053 | ( 0.856; 3.249) | **0.003** |
| BF |  | 0.026 | (0.024; 0.029) | **<2e-16** | * |  |  |  |  |  |
| BMI |  | 0.049 | (0.047; 0.052) | **<2e-16** | 339224 | 11 | 0.0022 | 0.351 | ( 0.118; 0.583) | **0.013** |
| WHR |  | 0.044 | (0.042; 0.047) | **<2e-16** | 210082 | 17 | 0.0022 | 0.220 | ( 0.052; 0.388) | **0.010** |
| **UAlb/UCr** |  |  |  |  |  |  |  |  |  |  |
| AF | 9251 | 1.155 | (1.130; 1.180) | **<2e-16** | 60620 | 37 | 0.0063 | 0.105 | ( -0.064; 0.274) | 0.232 |
| CAD | 7397 | 1.042 | (1.017; 1.067) | **6.87E-04** | * |  |  |  |  |  |
| HF | 2791 | 1.226 | (1.181; 1.272) | **<2e-16** | * |  |  |  |  |  |
| HS | 979 | 1.153 | (1.077; 1.233) | **3.90E-05** | * |  |  |  |  |  |
| IS | 1895 | 1.059 | (1.010; 1.111) | 0.018 | * |  |  |  |  |  |
| LIPID |  | 1.056 | (1.045; 1.068) | **<2e-16** | * |  |  |  |  |  |
| T2D |  | 1.243 | (1.222; 1.265) | **<2e-16** | 62892 | 25 | 0.0034 | 0.282 | ( 0.053; 0.511) | **0.023** |
| SBP |  | 0.157 | (0.154; 0.160) | **<2e-16** | 337222 | 37 | 0.0064 | 2.753 | ( 1.439; 4.068) | **2.06E-04** |
| **DBP** |  | 0.136 | (0.133; 0.139) | **<2e-16** | 337222 | 35 | 0.0064 | 0.848 | ( 0.078; 1.619) | **0.038** |
| **BF** |  | -0.106 | (-0.108; -0.103) | **<2e-16** | * |  |  |  |  |  |
| **BMI** |  | -0.170 | (-0.173; -0.167) | **<2e-16** | 339224 | 27 | 0.0039 | -0.056 | ( -0.184; 0.073) | 0.402 |
| **WHR** |  | -0.062 | (-0.064; -0.059) | **<2e-16** | 210082 | 29 | 0.0039 | 0.187 | ( 0.078; 0.295) | **7.36E-04** |

Main analyses were performed excluding participants with kidney disease and/or cardiovascular disease at baseline. Estimates are from multivariable-adjusted linear (continuous risk factors), Cox proportional hazards models (cardiovascular outcomes) or logistic (lipid treatment and T2D) regression, and inverse variance weighted method after outlier exclusion if needed for MR. The estimates represent SD change in outcome variable per SD change in the urinary biomarker tested. Binary outcomes: atrial fibrillation (AF), coronary artery disease (CAD), heart failure (HF), hemorrhagic stroke (HS), ischemic stroke (IS), lipid medications (LIPID) and type 2 diabetes (T2D). Continuous outcomes: systolic and diastolic blood pressure (SBP, and DBP; respectively), body fat percentage (BF), body mass index (BMI), waist-to-hip ratio (WHR). Models were adjusted for age, sex, region of the UK Biobank assessment center, ethnicity, smoking, alcohol, physical activity, Townsend index, blood pressure (DBP and SBP), obesity (BMI, body fat percentage, WHR), lipid medications, T2D, and medications affecting renal excretion. P-values significant after Bonferroni correction ( $p \leq 4.17E-03$ ) are bold. R<sup>2</sup>; proportion of phenotypic variance explained by the instruments (tested in UK Biobank).
MR analyses performed using publicly available consortia data, except for blood pressure where we performed a GWAS in UK Biobank (as the publicly available GWAS summary statistics were adjusted for BMI).
Abbreviations: HR, hazard ratio; OR, odds ratio; CI, confidence interval; SNP, single-nucleotide polymorphism; N, number of events. \* No MR analyses performed for lack of genome-wide association studies available or too low statistical power.

### Observational Analyses

Table 2 summarizes the results from our main observational analyses (full results in eTable 3). UNa/UK showed significant inverse associations (i.e. higher UNa/UK associated with lower disease risk) with AF, CAD, lipid-lowering medication and T2D. In contrast, higher UNa/UK was associated with higher SBP and DBP, as well as increased body fat percentage, BMI and WHR (*P*≤0.0042). When we performed these associations using sodium and potassium adjusted for creatinine, we detected similar and consistent results (eTable 3 and eFigure 1).

UAlb/UCr showed significant positive associations with AF, CAD, HF, HS, lipid-lowering medication and T2D. Further, high UAlb/UCr was associated with higher SBP and DBP. In contrast, UAlb/UCr showed significant inverse associations with body fat percentage, BMI, and WHR. The negative association with obesity traits (body fat percentage, BMI and WHR) was consistent also for UNa/UCr and UK/UCr (eTable 3) and was driven by the adjustment for creatinine. Indeed, urine sodium, potassium and albumin not adjusted for creatinine showed significant positive associations with obesity traits (eTable 4).

In our sensitivity analyses, after excluding participants using diuretics, ACE inhibitors, ARBs, or CCBs, we observed consistent results. For a few associations, UNa/UK with CAD, IS and T2D; UNa/UCr and UK/UCr with HF, and UAlb/UCr with CAD, we observed consistent directions with the main analyses, without reaching significance, probably due to the lower sample size (lower statistical power) (eTable 3 and eFigure 1).

We excluded nonlinear associations between UNa/UK and UAlb/UCr and all outcomes tested (*P*>0.05), except for HF (*P*=9.0E10-8, UNa/UK) and IS (*P*=0.001, UAlb/UCr) by spline regression (eFigure 2 and 3).

When participants eligible for inclusion in the main analysis were stratified by sex (eTable 5 and eFigure 4), no additional significant associations were found between exposures and outcomes in either subset. All associations between urinary biomarkers and outcomes remained significant and consistent with the main analyses in both men and women. Events were more common in the male sample set for all CVD outcomes (eTable 5). Generally, males displayed larger effect estimates than females. For disease outcomes, this potentially could be explained by better statistical power (more events), while the power should be equal for continuous traits (as the number of measurements were similar). The strongest sex interactions were observed for UAlb/UCr and T2D (P<2E10-16), and across urinary biomarkers and obesity traits (P<2E-16) (eTable 5 and eFigure 4).

### Mendelian Randomization

After correcting for horizontal pleiotropy, we found evidence of causal associations between UNa/UK and BMI, UAlb/UCr and T2D, and of both biomarkers (UNa/UK and UAlb/UCr) with blood pressure and WHR (Table 2 and Figure 1). A leave-one out sensitivity analysis did not highlight any SNPs with a large effect on the results. After excluding heterogeneous SNPs using MR-PRESSO (eFigures 5-8), our analysis showed no significant heterogeneity and no significant directional horizontal pleiotropy. Numbers of variants included in the analyses, number of outliers excluded, and full results are shown in eTable 6. We only performed MR analyses for outcomes for which we had at least 75% statistical power (eTable 2).

**Figure 1.**
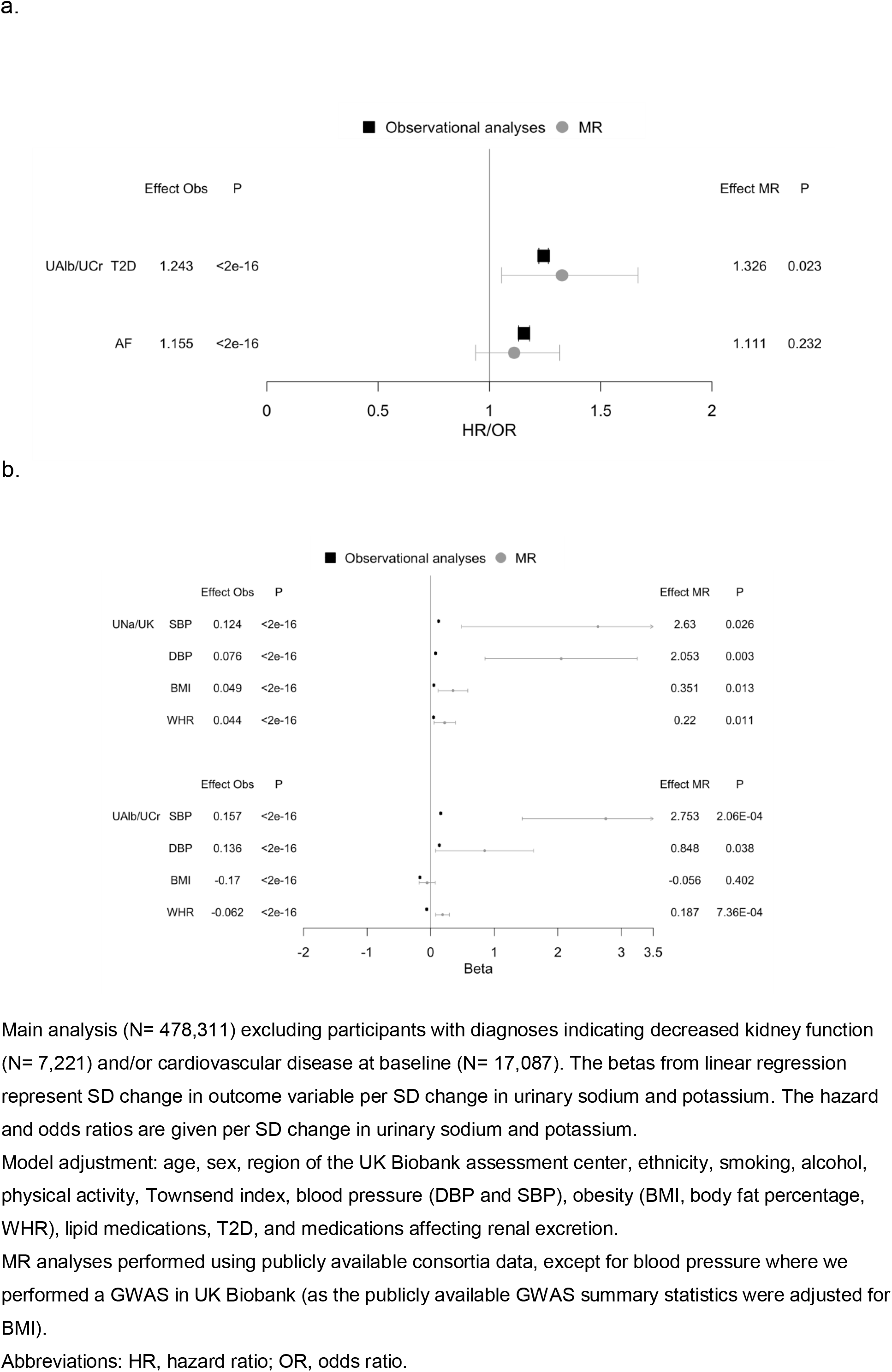
Observational and Mendelian randomization (MR) analyses of urinary sodium-potassium ratio (UNa/UK) and urinary albumin-creatinine ratio (UAlb/UCr) with cardiovascular outcomes in UK Biobank using multivariable-adjusted linear, logistic and Cox proportional hazards models in main observational analyses and inverse variance weighted method. a. Associations of UAlb/UCr with type 2 diabetes (T2D) and atrial fibrillation (AF); b. Associations of UNa/UK and UAlb/UCr with systolic and diastolic blood pressure (SBP and DBP), body mass index (BMI), and waist-to-hip ratio (WHR).

We found evidence of causal bidirectional effect across UNa/UK and UAlb/UCr and blood pressure, and between albumin and T2D (eTable 7).

We performed multivariable MR using all established GWAS significant variants for UAlb/UCr, SBP and WHR as predictor variables, and GWAS of T2D and AF as outcome variables. We detected an independent association between UAlb/UCr and T2D, while the association of UAlb/UCr with AF was mediated by SBP (eTable 8).

## Discussion

### Principal Findings

We studied associations of urinary biomarkers with cardiometabolic disease in 478,311 individuals free of chronic kidney disease and CVD at baseline. We made four main findings. First, we found a positive association between UNa/UK and albumin with blood pressure, as well as with adiposity-related measures (body fat percentage, BMI, or WHR). Second, we observed a direct association of UAlb/UCr with CVD incidence, but an inverse association of UNa/UK with incident CVD and T2D in traditional observational analyses. Third, the strongest sex differences were observed for associations between UAlb/UCr and T2D, and across urinary biomarkers and obesity traits. However, all associations were directionally consistent. Lastly, using a Mendelian randomization approach, we provide evidence that higher UNa/UK is causally related with higher blood pressure, and we highlight a feed-back causal loop between albumin and hypertension, and between albumin and T2D.

### Comparison with Prior Observational Studies

Our results are consistent with previous literature which reported that high albumin is associated with higher susceptibility to CVD^13^ and hypertension.^15^ Previous studies have shown inconsistent results between sodium intake and the risk of CVD. The Scottish Heart Study, a population-based longitudinal study of 10,000 people, showed no consistent associations between sodium intake and cardiovascular or all-cause mortality.^23^ On the other hand, previous meta-analyses of observational studies have suggested a direct association of sodium intake with stroke and CAD events,^12^ and a potential inverse association between potassium intake and risk of stroke and other CVD outcomes.^24^ In agreement with these findings, randomized interventional trials have shown that sodium reduction can reduce the long term risk of developing high blood pressure,^25^ cardiovascular events^8,9^ and death.^10^ Our findings from observational analyses show an inverse relationship between UNa/UK excretion and AF, CAD, and IS. A study recently published in the *Lancet* found a positive association between sodium intake and stroke only in communities where mean intake was greater than 5 g/day, largely confined to China. By contrast, they found an inverse relation with myocardial infarction and mortality. The same study also showed that the rates of stroke, cardiovascular death, and total mortality decreased with increasing potassium intake across all the communities studied.^26^

We detected an inverse association between UNa/UK and hyperlipidemia in line with a previous meta-analysis of randomized controlled trials which found that a low sodium diet intervention was associated with an increase in cholesterol and triglycerides.^27^ We also identified an inverse association between UNa/UK and T2D in the UK Biobank cohort. To the best of our knowledge, the relationship between sodium intake and incidence of T2D has not been previously described. In contrast, the association between albumin and diabetes was already observed in previous studies^28^, and it is consistent with the positive association detected by our study.

We also confirmed the well-established direct association between sodium intake and blood pressure, as well as the inverse association between potassium intake and blood pressure^4–7^ in the UK Biobank cohort. In addition, we replicated these associations using urinary sodium and urinary potassium adjusted for creatinine. Regarding albumin and blood pressure, our results are consistent with a recent previous study^15^ which supported the existence of a bidirectional causal association between albuminuria and blood pressure.

We observed positive associations between UNa/UK and albumin with obesity and adiposity-related risk factors (body fat percentage, BMI, WHR). These results are consistent with previous studies that have suggested that sodium intake is an independent risk factor for obesity,^29,30^ and that albuminuria is one the most important risk factors for obesity^31^.

No previous MR studies have investigated the causality of UNa/UK with cardiovascular outcomes and risk factors, and of albumin with T2D. Our MR results mirror and extend findings from previous randomized interventional trials^4,5^ that have established sodium intake as a risk factor for hypertension. In addition, we highlight a feed-back causal loop between albumin and T2D, and we also replicate the existence of a bidirectional causal association between albuminuria and blood pressure that was reported by a recent MR study.^15^

### The Potential Causal Role of Urinary Biomarkers in Blood Pressure and Type 2 Diabetes Mellitus

The incidence and prevalence of hypertension continue to rise, presumably due to an aging population, increasing obesity and physical inactivity.^32^ Substantial evidence from clinical trials have demonstrated that high sodium and low potassium intake are significantly associated with elevated blood pressure.^4,5^ Previous studies also show that reducing dietary sodium in individuals with prehypertension decrease the risk of cardiovascular events and overall mortality.^8–10^ This is the first study exploring this association using genetic data by means of a MR approach. We confirm a causal role of high sodium and low potassium excretion – arguably reflecting dietary intake – in higher susceptibility for increased blood pressure. A high-sodium diet suppresses the renin-angiotensin-aldosterone system. Endothelial dysfunction probably plays an important role to modulate the influence of high sodium intake on blood pressure, although the exact mechanisms remain elusive.^33^ We also observed a bidirectional causal association of albumin with blood pressure. The causal association of albumin with blood pressure was already highlighted in a recent study that suggested the existence of a feed-forward loop where elevated blood pressure leads to increased albuminuria, which in turn further increases blood pressure.^15^ The presence of albuminuria is a powerful predictor of renal and cardiovascular risk in patients with T2D and hypertension. Multiple studies have shown that decreasing albuminuria reduces the risk of adverse renal and cardiovascular outcomes. The pathophysiology is not definitively known, but also in this case, it is hypothesized to be related to endothelial dysfunction, inflammation, and/or abnormalities in the renin-angiotensin-aldosterone system.^34^

Glomerular endothelial dysfunction is also implicated in the link between albuminuria and T2D.^35,36^ Our data suggest an independent causal association between T2D and albumin, not mediated by blood pressure. Urine albumin arises primarily from the increased passage of albumin through the glomerular filtration barrier that is insufficiently reabsorbed by tubular epithelium. The filtration barrier is comprised of the endothelium, glomerular basement membrane, and podocytes. T2D can lead to disruption of each of these components including the endothelium in which disruption of the endothelial glycocalyx through dysregulation by the diabetic milieu.^35^ Indeed, diabetic patients have decreased systemic glycocalyx volume, and this is correlated with the presence of albuminuria.^37^

### Strengths and Limitations

Our study is the largest and most comprehensive study of associations of urinary biomarkers with cardiovascular risk factors, T2D and CVD to date. Strengths of our study include the large sample size, the robustness of our findings, the most recent and powerful GWAS summary statistics as outcomes, and a new method, the MR-PRESSO, to decrease the risk of pleiotropy.

Our study also has several limitations. First, we are limited to using measures currently available in the UK Biobank. We excluded participants with diagnoses indicating kidney disease, but using an estimated glomerular filtration rate would have been a more accurate method to exclude individuals with mild CKD. Such an exclusion cannot be performed until serum creatinine measurements are available in the UK Biobank in the future. Second, the vast majority of participants were of European ancestry despite the inclusion of several non-European ethnicities. Hence, our results may not be generalizable to other race/ethnic groups with significantly different diets and/or prevalence and predispositions to cardiometabolic disease. Finally, statistical power to detect potentially causal relationships through our MR studies was limited for some traits, at least for smaller effects, including some of those observed in our traditional epidemiological analyses.

## Conclusions

Our study mirrors and extends findings from randomized interventional trials which have established sodium intake as a risk factor for hypertension. In addition, we detect a feed-back causal loop between albumin and hypertension, and our finding of a bidirectional causal association between albumin and T2D reflects the well-known nephropathy in T2D.

## Acknowledgements

This research has been conducted using the UK Biobank Resource under Application Number 13721. The research was performed with support from National Institutes of Health (1R01HL135313-01; 1R01DK106236-01A1) and the Stanford Diabetes Research center award (P30DK116074).

