## Supplemental Material for "Urinary biomarkers and cardiovascular outcomes in the UK Biobank: observational and Mendelian randomization analyses"

### Supplemental Tables

**eTable 1.** Variants included in the instrument variable for Mendelian randomization analyses.

| **UNa/UK** | | | | | | |
|---|---|---|---|---|---|---|
| **SNP** | **EA** | **OA** | **EAF** | **BETA** | **SE** | **P** |
| rs1194277 | G | C | 0.508 | 0.015 | 0.002 | 1.57E-10 |
| rs7157772 | T | G | 0.612 | 0.015 | 0.002 | 4.66E-10 |
| rs299938 | G | A | 0.234 | -0.017 | 0.003 | 2.13E-09 |
| rs8096658 | G | C | 0.488 | -0.014 | 0.002 | 2.69E-08 |
| rs2903625 | T | A | 0.384 | 0.014 | 0.002 | 2.41E-08 |
| rs34783010 | T | G | 0.193 | -0.022 | 0.003 | 7.07E-13 |
| rs1698114 | G | A | 0.484 | 0.016 | 0.003 | 1.39E-09 |
| rs6754311 | C | T | 0.248 | 0.017 | 0.003 | 1.03E-09 |
| rs3770435 | C | T | 0.362 | 0.017 | 0.003 | 3.06E-11 |
| rs4665972 | C | T | 0.607 | -0.024 | 0.002 | 2.89E-22 |
| rs11373337 | NA | A | 0.705 | 0.015 | 0.003 | 6.74E-09 |
| rs6440008 | C | T | 0.386 | -0.017 | 0.002 | 4.07E-12 |
| rs12496573 | A | G | 0.211 | 0.016 | 0.003 | 3.36E-08 |
| rs2035562 | G | A | 0.672 | -0.014 | 0.003 | 4.02E-08 |
| rs1229984 | C | T | 0.978 | -0.056 | 0.008 | 3.82E-12 |
| rs9291156 | G | A | 0.455 | -0.014 | 0.002 | 7.09E-09 |
| rs2057655 | A | G | 0.185 | -0.017 | 0.003 | 2.31E-08 |
| rs13188076 | T | G | 0.232 | -0.020 | 0.003 | 2.22E-12 |
| rs56108664 | T | C | 0.199 | 0.020 | 0.003 | 4.29E-11 |
| rs34867029 | T | NA | 0.704 | -0.015 | 0.003 | 3.94E-08 |
| rs2455357 | G | A | 0.705 | 0.016 | 0.003 | 4.25E-08 |
| rs6452788 | A | G | 0.236 | -0.021 | 0.003 | 1.79E-13 |
| rs2394893 | A | G | 0.605 | 0.014 | 0.002 | 2.27E-08 |
| rs791903 | C | G | 0.556 | 0.013 | 0.002 | 2.72E-08 |
| rs62458486 | A | G | 0.226 | -0.016 | 0.003 | 2.37E-08 |
| rs372106593 | NA | G | 0.446 | -0.013 | 0.002 | 4.30E-08 |
| rs11514731 | G | C | 0.179 | -0.020 | 0.003 | 1.28E-10 |
| rs12700027 | A | G | 0.110 | 0.023 | 0.004 | 2.07E-09 |
| rs33951980 | T | C | 0.130 | -0.021 | 0.004 | 7.08E-09 |
| rs4873492 | T | C | 0.170 | 0.020 | 0.003 | 6.29E-10 |
| rs11103388 | A | G | 0.330 | 0.014 | 0.003 | 2.75E-08 |

| UNa/UCr | | | | | | |
|---|---|---|---|---|---|---|
| **SNP** | **EA** | **OA** | **EAF** | **BETA** | **SE** | **P** |
| rs12363886 | A | C | 0.482 | -0.013 | 0.002 | 3.29E-08 |
| rs11062590 | G | C | 0.092 | -0.025 | 0.004 | 2.42E-09 |
| rs35335867 | NA | C | 0.248 | -0.019 | 0.003 | 1.15E-11 |
| rs11659764 | A | T | 0.054 | 0.030 | 0.005 | 3.08E-08 |
| rs8096658 | G | C | 0.488 | -0.014 | 0.002 | 7.96E-09 |
| rs1698114 | G | A | 0.484 | 0.015 | 0.003 | 9.36E-09 |
| rs1047891 | A | C | 0.316 | -0.027 | 0.003 | 1.34E-24 |
| rs35169442 | C | G | 0.198 | 0.021 | 0.003 | 9.87E-12 |
| rs1260326 | C | T | 0.607 | -0.028 | 0.002 | 4.05E-29 |
| rs62140396 | A | G | 0.118 | 0.022 | 0.004 | 5.33E-09 |
| rs13056137 | A | C | 0.264 | 0.017 | 0.003 | 2.53E-10 |
| rs6440008 | C | T | 0.386 | -0.017 | 0.003 | 4.65E-12 |
| rs2035561 | C | T | 0.671 | -0.014 | 0.003 | 3.29E-08 |
| rs1229984 | C | T | 0.978 | -0.050 | 0.008 | 7.36E-10 |
| rs13127170 | G | A | 0.552 | 0.014 | 0.002 | 1.61E-08 |
| rs4697700 | C | G | 0.236 | 0.016 | 0.003 | 1.17E-08 |
| rs13188076 | T | G | 0.232 | -0.024 | 0.003 | 6.48E-17 |
| rs1423553 | C | T | 0.672 | 0.014 | 0.003 | 3.75E-08 |
| rs702634 | A | G | 0.692 | 0.016 | 0.003 | 6.62E-10 |
| rs13242739 | A | T | 0.051 | -0.031 | 0.006 | 2.37E-08 |
| rs33951980 | T | C | 0.130 | -0.023 | 0.004 | 1.44E-10 |
| rs2953661 | C | A | 0.696 | -0.017 | 0.003 | 4.02E-09 |
| rs2954021 | G | A | 0.506 | -0.016 | 0.002 | 1.36E-10 |
| rs558455 | G | A | 0.580 | 0.016 | 0.002 | 1.67E-10 |
| rs4873492 | T | C | 0.170 | 0.022 | 0.003 | 1.07E-11 |
| rs12378270 | A | G | 0.061 | 0.032 | 0.006 | 6.49E-09 |
| **UK/UCr** | | | | | | |
| **SNP** | **EA** | **OA** | **EAF** | **BETA** | **SE** | **P** |
| rs35041900 | T | C | 0.093 | -0.024 | 0.004 | 1.74E-09 |
| rs66495454 | NA | G | 0.378 | 0.014 | 0.002 | 9.63E-09 |
| rs2664284 | C | G | 0.255 | 0.018 | 0.003 | 5.84E-12 |
| rs4408310 | A | C | 0.539 | 0.014 | 0.002 | 4.33E-09 |
| rs57739486 | T | C | 0.158 | 0.017 | 0.003 | 2.75E-08 |
| rs35335867 | NA | C | 0.248 | -0.024 | 0.003 | 8.42E-19 |
| rs2472297 | T | C | 0.267 | 0.021 | 0.003 | 4.44E-16 |
| rs11648252 | G | C | 0.795 | 0.016 | 0.003 | 2.09E-08 |

| | | | | | | |
|---|---|---|---|---|---|---|
| rs71352040 | A | T | 0.071 | 0.025 | 0.004 | 1.17E-08 |
| rs4803380 | T | C | 0.022 | -0.047 | 0.008 | 1.53E-09 |
| rs34783010 | T | G | 0.193 | 0.026 | 0.003 | 3.10E-19 |
| rs2603274 | C | A | 0.817 | 0.018 | 0.003 | 1.53E-09 |
| rs1047891 | A | C | 0.316 | -0.031 | 0.002 | 1.60E-37 |
| rs4973766 | T | C | 0.456 | -0.022 | 0.002 | 4.29E-21 |
| rs1664781 | A | G | 0.693 | 0.014 | 0.002 | 1.75E-08 |
| rs72704785 | A | G | 0.179 | -0.024 | 0.003 | 1.03E-14 |
| rs10872423 | T | C | 0.194 | -0.019 | 0.003 | 2.46E-11 |
| rs3129996 | C | A | 0.906 | -0.021 | 0.004 | 4.20E-08 |
| rs2260051 | T | A | 0.558 | -0.013 | 0.002 | 1.44E-08 |
| rs9269041 | A | G | 0.647 | -0.014 | 0.002 | 1.89E-09 |
| rs77213792 | G | C | 0.068 | -0.027 | 0.005 | 2.78E-09 |
| rs10950655 | C | T | 0.505 | 0.013 | 0.002 | 4.53E-09 |
| rs756018 | G | C | 0.110 | -0.026 | 0.004 | 1.19E-12 |
| **UAlb/UCr** | | | | | | |
| **SNP** | **EA** | **OA** | **EAF** | **BETA** | **SE** | **P** |
| rs35388680 | NA | C | 0.407 | 0.013 | 0.002 | 1.92E-08 |
| rs35202981 | G | A | 0.140 | 0.020 | 0.003 | 3.03E-09 |
| rs58953262 | NA | C | 0.521 | 0.013 | 0.002 | 4.48E-08 |
| rs1204672 | C | T | 0.572 | 0.014 | 0.002 | 5.35E-09 |
| rs12032996 | A | G | 0.162 | -0.018 | 0.003 | 1.59E-08 |
| rs17358122 | T | C | 0.220 | -0.015 | 0.003 | 3.71E-08 |
| rs55668525 | T | G | 0.192 | 0.017 | 0.003 | 4.03E-09 |
| rs10157710 | T | C | 0.801 | 0.025 | 0.003 | 4.48E-18 |
| rs3781470 | A | C | 0.355 | 0.013 | 0.002 | 3.11E-08 |
| rs79051478 | C | G | 0.104 | 0.058 | 0.004 | 1.50E-52 |
| rs780836 | C | G | 0.715 | 0.014 | 0.003 | 2.56E-08 |
| rs188126412 | T | C | 0.028 | 0.073 | 0.007 | 6.58E-24 |
| rs116867125 | A | G | 0.019 | 0.059 | 0.009 | 6.08E-12 |
| rs79386054 | T | C | 0.017 | 0.055 | 0.010 | 9.79E-09 |
| rs7910002 | C | G | 0.698 | 0.014 | 0.003 | 2.45E-08 |
| rs67339103 | A | G | 0.212 | 0.020 | 0.003 | 7.90E-12 |
| rs7115200 | G | T | 0.440 | 0.013 | 0.002 | 1.14E-08 |
| rs34183365 | NA | C | 0.415 | 0.014 | 0.002 | 3.73E-08 |
| rs2601006 | T | C | 0.343 | -0.014 | 0.002 | 2.14E-08 |
| rs8036643 | C | G | 0.645 | 0.015 | 0.002 | 8.72E-10 |
| rs35335867 | NA | C | 0.248 | -0.019 | 0.003 | 1.33E-12 |

| | | | | | | |
|---|---|---|---|---|---|---|
| rs146311723 | C | T | 0.174 | 0.017 | 0.003 | 2.19E-08 |
| rs2472297 | T | C | 0.267 | 0.025 | 0.003 | 2.65E-21 |
| rs9673084 | A | G | 0.271 | 0.019 | 0.003 | 3.06E-13 |
| rs4488444 | G | A | 0.756 | -0.016 | 0.003 | 7.55E-09 |
| rs11078597 | C | T | 0.186 | 0.017 | 0.003 | 1.87E-08 |
| rs35572189 | A | G | 0.361 | -0.013 | 0.002 | 3.82E-08 |
| rs784257 | C | T | 0.813 | -0.017 | 0.003 | 5.10E-09 |
| rs11882796 | T | A | 0.542 | 0.014 | 0.002 | 3.58E-09 |
| rs10419198 | T | C | 0.250 | -0.015 | 0.003 | 4.20E-08 |
| rs10207567 | C | G | 0.812 | 0.020 | 0.003 | 3.51E-11 |
| rs1047891 | A | C | 0.316 | -0.018 | 0.002 | 8.39E-13 |
| rs35687319 | T | C | 0.128 | 0.022 | 0.004 | 4.95E-10 |
| rs4665972 | C | T | 0.607 | -0.017 | 0.002 | 4.31E-13 |
| rs2627770 | T | C | 0.732 | -0.015 | 0.003 | 1.01E-08 |
| rs12714144 | T | A | 0.127 | -0.021 | 0.003 | 1.91E-09 |
| rs2295762 | G | A | 0.360 | 0.014 | 0.002 | 2.11E-08 |
| rs112607182 | T | C | 0.075 | 0.029 | 0.005 | 3.55E-10 |
| rs6535594 | A | G | 0.495 | 0.013 | 0.002 | 1.17E-08 |
| rs4109437 | A | G | 0.038 | 0.057 | 0.006 | 2.58E-21 |
| rs146996008 | NA | A | 0.419 | 0.014 | 0.002 | 5.57E-09 |
| rs13179493 | C | T | 0.293 | -0.014 | 0.003 | 3.09E-08 |
| rs114118094 | T | A | 0.075 | -0.030 | 0.004 | 8.96E-12 |
| rs40480 | G | C | 0.362 | 0.014 | 0.002 | 2.75E-09 |
| rs10947788 | T | C | 0.289 | 0.015 | 0.003 | 1.41E-08 |
| rs4410790 | C | T | 0.634 | 0.024 | 0.002 | 1.00E-22 |
| rs3735533 | C | T | 0.926 | 0.025 | 0.004 | 2.36E-08 |
| rs17158386 | A | G | 0.261 | 0.019 | 0.003 | 2.45E-12 |
| rs28601761 | G | C | 0.420 | -0.017 | 0.002 | 4.03E-13 |

Abbreviations: UNa/UK, urinary sodium to potassium ratio; UNa/UCr, urinary sodium to creatinine ratio; UK/UCr, urinary potassium to creatinine ratio; UAlb/UCr, urinary albumin to creatinine ratio; SNP, single-nucleotide polymorphism; EA, effect allele; OA, other allele; EAF, effect allele frequency; SE, standard error; P, p value.

**eTable 2.** Description of data used and statistical power for Mendelian randomization.

| | | | UNa/UK | | | | UNa/UCr | | | | UK/UCr | | | | UAlb/UCr | | | |
|---|---|---|---|---|---|---|---|---|---|---|---|---|---|---|---|---|---|---|
| **Outcomes** | **PMID** | **Sample size** | **R2*** | **Effect obs** | **Power obs effect** | **Power fixed effect** | **R2*** | **Effect obs** | **Power obs effect** | **Power fixed effect** | **R2*** | **Effect obs** | **Power obs effect** | **Power fixed effect** | **R2*** | **Effect obs** | **Power obs effect** | **Power fixed effect** |
| **AF** | **30061737** | 1,030,836 | 0.0031 | 0.898 | 30 | 46 | 0.0031 | 0.905 | 26 | 46 | 0.0031 | 0.986 | 4 | 46 | 0.0063 | 1.155 | **78** | **76** |
| **CAD** | **26343387** | 184,305 | 0.0032 | 0.962 | 7 | 36 | 0.0032 | 0.936 | 11 | 36 | 0.0028 | 0.947 | 8 | 32 | 0.0060 | 1.042 | 9 | 59 |
| **Stroke** | **29531354** | 446,696 | 0.0031 | 0.929 | 12 | 32 | 0.0033 | 0.879 | 30 | 34 | 0.0030 | 0.892 | 22 | 31 | 0.0063 | 1.059 | 14 | 57 |
| **T2D** | **30054458** | 659,316 | 0.0018 | 0.968 | 5 | 30 | 0.0025 | 0.990 | 3 | 40 | 0.0018 | 1.036 | 6 | 30 | 0.0034 | 1.243 | **88** | 51 |
| **SBP** | | 337,222 | 0.0034 | 0.124 | **99** | **100** | 0.0033 | 0.120 | **98** | **100** | 0.0031 | -0.009 | 5 | **100** | 0.0064 | 0.157 | **100** | **100** |
| **DBP** | | 337,222 | 0.0034 | 0.076 | 73 | **100** | 0.0033 | 0.078 | 74 | **100** | 0.0031 | 0.001 | 3 | **100** | 0.0064 | 0.136 | **100** | **100** |
| **SBP** | **28739976** | 150,134 | 0.0028 | 0.124 | 72 | **87** | 0.0029 | 0.120 | 70 | **88** | 0.0027 | -0.009 | 4 | **86** | 0.0055 | 0.157 | **100** | **99** |
| **SBP** | **28739976** | 150,134 | 0.0028 | 0.076 | 35 | **87** | 0.0029 | 0.078 | 37 | **88** | 0.0027 | 0.001 | 3 | **86** | 0.0055 | 0.136 | **98** | **99** |
| **BMI** | **25673413** | 339,224 | 0.0022 | 0.049 | 26 | **98** | 0.0021 | -0.054 | 31 | **98** | 0.0019 | -0.154 | **98** | **98** | 0.0039 | -0.170 | **100** | **100** |
| **WHR** | **25673412** | 210,082 | 0.0022 | 0.044 | 15 | **90** | 0.0021 | -0.007 | 4 | **88** | 0.0019 | -0.076 | 34 | **85** | 0.0039 | -0.062 | 42 | **99** |

Characteristics of the consortia used in our study: number of samples, proportion of phenotypic variance explained by the instruments (tested in UK Biobank), statistical power considering the effect from observational analyses and for a fixed effect of 1.15 (binary traits) and 0.15 (continuous traits). Power estimates >75% in bold.

Abbreviations: effect obs, effect from observational analyses; AF, atrial fibrillation; CAD, coronary artery disease; T2D, type 2 diabetes; SBP and DBP, systolic and diastolic blood pressure, respectively; BMI, body mass index; WHR, waist-to-hip ratio.

* The variance explained varied somewhat across different outcomes due to lack of variants in some GWAS summary statistics.

**eTable 3.** Observational associations of the urinary sodium to potassium ratio (UNa/UK), sodium and potassium adjusted for creatinine (UNa/UCr and UK/UCr), and urinary albumin to creatinine ratio (UAlb/UCr) with cardiovascular outcomes in UK Biobank using multivariable-adjusted Cox proportional hazards models, and multivariable-adjusted linear and logistic regression for main and sensitivity analyses.

| **UNa/UK** | **Main** | | | | | **Sensitivity** | | | | |
|---|---|---|---|---|---|---|---|---|---|---|
| **Outcome** | **N** | **HR/OR/Beta** | **Lower** | **Upper** | **P** | **N** | **HR/OR/Beta** | **Lower** | **Upper** | **P** |
| **AF** | 9,196 | 0.898 | 0.879 | 0.917 | **<2e-16** | 5,499 | 0.895 | 0.869 | 0.921 | **5.03E-14** |
| **CAD** | 7,375 | 0.962 | 0.939 | 0.985 | **1.24E-03** | 4,659 | 0.977 | 0.947 | 1.009 | 1.53E-01 |
| **HF** | 2,775 | 0.960 | 0.924 | 0.997 | 3.41E-02 | 1,422 | 0.999 | 0.944 | 1.057 | 9.72E-01 |
| **HS** | 978 | 0.982 | 0.92 | 1.049 | 5.91E-01 | 714 | 0.967 | 0.893 | 1.047 | 4.07E-01 |
| **IS** | 1,888 | 0.929 | 0.886 | 0.973 | **1.94E-03** | 1,268 | 0.946 | 0.891 | 1.004 | 6.69E-02 |
| **LIPID** | 67,898 | 0.940 | 0.93 | 0.957 | **<2e-16** | 28,978 | 0.913 | 0.899 | 0.926 | **<2e-16** |
| **T2D** | 19,876 | 0.968 | 0.951 | 0.984 | **1.84E-04** | 7,758 | 0.986 | 0.959 | 1.014 | 3.16E-01 |
| **SBP** | | 0.124 | 0.121 | 0.127 | **<2e-16** | | 0.131 | 0.127 | 0.134 | **<2e-16** |
| **DBP** | | 0.076 | 0.073 | 0.079 | **<2e-16** | | 0.080 | 0.077 | 0.083 | **<2e-16** |
| **BF** | | 0.026 | 0.024 | 0.029 | **<2e-16** | | 0.022 | 0.019 | 0.024 | **<2e-16** |
| **BMI** | | 0.049 | 0.047 | 0.052 | **<2e-16** | | 0.049 | 0.046 | 0.052 | **<2e-16** |
| **WHR** | | 0.044 | 0.042 | 0.047 | **<2e-16** | | 0.045 | 0.042 | 0.048 | **<2e-16** |
| **UNa/UCr** | **Main** | | | | | **Sensitivity** | | | | |
| **Outcome** | **N** | **HR/OR/Beta** | **Lower** | **Upper** | **P** | **N** | **HR/OR/Beta** | **Lower** | **Upper** | **P** |
| **AF** | 9,219 | 0.905 | 0.887 | 0.924 | **<2e-16** | 5,516 | 0.895 | 0.87 | 0.922 | **8.48E-14** |
| **CAD** | 7,387 | 0.936 | 0.914 | 0.958 | **1.93E-08** | 4,665 | 0.942 | 0.913 | 0.973 | **2.37E-04** |
| **HF** | 2,776 | 0.937 | 0.904 | 0.972 | **4.76E-04** | 1,423 | 0.967 | 0.914 | 1.024 | 2.47E-01 |
| **HS** | 978 | 0.955 | 0.897 | 1.018 | 1.60E-01 | 714 | 0.965 | 0.892 | 1.045 | 3.82E-01 |

| | | | | | | | | | | |
|---|---|---|---|---|---|---|---|---|---|---|
| **IS** | 1,888 | 0.879 | 0.84 | 0.92 | **3.04E-08** | 1,268 | 0.881 | 0.83 | 0.935 | **3.26E-05** |
| **LIPID** | 67,898 | 0.958 | 0.948 | 0.967 | **<2e-16** | 28,978 | 0.942 | 0.928 | 0.956 | **2.46E-15** |
| **T2D** | 19,876 | 0.990 | 0.974 | 1.006 | 2.25E-01 | 7,758 | 1.024 | 0.995 | 1.053 | 1.02E-01 |
| **SBP** | | 0.120 | 0.118 | 0.123 | **<2e-16** | | 0.128 | 0.125 | 0.131 | **<2e-16** |
| **DBP** | | 0.078 | 0.076 | 0.081 | **<2e-16** | | 0.084 | 0.08 | 0.087 | **<2e-16** |
| **BF** | | -0.040 | -0.042 | -0.037 | **<2e-16** | | -0.048 | -0.05 | -0.045 | **<2e-16** |
| **BMI** | | -0.054 | -0.057 | -0.051 | **<2e-16** | | -0.061 | -0.064 | -0.058 | **<2e-16** |
| **WHR** | | -0.007 | -0.009 | -0.005 | **1.58E-09** | | -0.009 | -0.011 | -0.006 | **2.08E-11** |
| **UK/UCr** | **Main** | | | | | **Sensitivity** | | | | |
| **Outcome** | **N** | **HR/OR/Beta** | **Lower** | **Upper** | **P** | **N** | **HR/OR/Beta** | **Lower** | **Upper** | **P** |
| **AF** | 9,229 | 0.986 | 0.962 | 1.009 | 2.35E-01 | 5,513 | 0.997 | 0.967 | 1.028 | 8.50E-01 |
| **CAD** | 7,385 | 0.947 | 0.922 | 0.972 | **5.46E-05** | 4,666 | 0.945 | 0.914 | 0.977 | **9.22E-04** |
| **HF** | 2,789 | 0.916 | 0.877 | 0.956 | **6.07E-05** | 1,425 | 0.934 | 0.879 | 0.992 | 2.52E-02 |
| **HS** | 979 | 0.942 | 0.879 | 1.011 | 9.83E-02 | 715 | 0.984 | 0.907 | 1.067 | 6.95E-01 |
| **IS** | 1,894 | 0.892 | 0.847 | 0.939 | **1.57E-05** | 1,270 | 0.890 | 0.836 | 0.948 | **2.89E-04** |
| **LIPID** | 67,898 | 1.030 | 1.018 | 1.043 | **6.52E-07** | 28,978 | 1.050 | 1.034 | 1.067 | **8.47E-10** |
| **T2D** | 19,876 | 1.036 | 1.016 | 1.057 | **4.82E-04** | 7,758 | 1.051 | 1.02 | 1.083 | **1.10E-03** |
| **SBP** | | -0.009 | -0.012 | -0.006 | **2.80E-09** | | -0.015 | -0.018 | -0.012 | **<2e-16** |
| **DBP** | | 0.001 | -0.002 | 0.004 | 5.45E-01 | | -0.003 | -0.007 | 0 | 4.56E-02 |
| **BF** | | -0.099 | -0.101 | -0.096 | **<2e-16** | | -0.095 | -0.098 | -0.093 | **<2e-16** |
| **BMI** | | -0.154 | -0.157 | -0.151 | **<2e-16** | | -0.150 | -0.153 | -0.147 | **<2e-16** |
| **WHR** | | -0.076 | -0.079 | -0.074 | **<2e-16** | | -0.074 | -0.077 | -0.071 | **<2e-16** |
| **UAlb/UCr** | **Main** | | | | | **Sensitivity** | | | | |
| **Outcome** | **N** | **HR/OR/Beta** | **Lower** | **Upper** | **P** | **N** | **HR/OR/Beta** | **Lower** | **Upper** | **P** |
| **AF** | 9,251 | 1.155 | 1.13 | 1.18 | **<2e-16** | 5,529 | 1.113 | 1.081 | 1.146 | **4.40E-13** |

| | | | | | | | | | | |
|---|---|---|---|---|---|---|---|---|---|---|
| **CAD** | 7,397 | 1.042 | 1.017 | 1.067 | **6.87E-04** | 4,672 | 1.015 | 0.984 | 1.047 | 3.56E-01 |
| **HF** | 2,791 | 1.226 | 1.181 | 1.272 | **<2e-16** | 1,425 | 1.210 | 1.145 | 1.28 | **1.61E-11** |
| **HS** | 979 | 1.153 | 1.077 | 1.233 | **3.90E-05** | 715 | 1.132 | 1.043 | 1.228 | **2.95E-03** |
| **IS** | 1,895 | 1.059 | 1.01 | 1.111 | 1.82E-02 | 1,271 | 1.031 | 0.971 | 1.095 | 3.22E-01 |
| **LIPID** | 67,898 | 1.056 | 1.045 | 1.068 | **<2e-16** | 28,978 | 1.060 | 1.044 | 1.076 | **2.61E-14** |
| **T2D** | 19,876 | 1.243 | 1.222 | 1.265 | **<2e-16** | 7,758 | 1.226 | 1.193 | 1.26 | **<2e-16** |
| **SBP** | | 0.157 | 0.154 | 0.16 | **<2e-16** | | 0.161 | 0.157 | 0.164 | **<2e-16** |
| **DBP** | | 0.136 | 0.133 | 0.139 | **<2e-16** | | 0.145 | 0.141 | 0.148 | **<2e-16** |
| **BF** | | -0.106 | -0.108 | -0.103 | **<2e-16** | | -0.114 | -0.117 | -0.112 | **<2e-16** |
| **BMI** | | -0.170 | -0.173 | -0.167 | **<2e-16** | | -0.185 | -0.189 | -0.182 | **<2e-16** |
| **WHR** | | -0.062 | -0.064 | -0.059 | **<2e-16** | | -0.071 | -0.074 | -0.068 | **<2e-16** |

Main analyses were performed excluding participants with kidney disease and/or cardiovascular disease at baseline. Sensitivity analyses was performed excluding participants with kidney disease and/or cardiovascular disease at baseline and/or participants using medications affecting renal excretion. Estimates are from multivariable-adjusted linear (continuous risk factors), Cox proportional hazards models (cardiovascular outcomes) or logistic (lipid treatment and T2D) regression. The estimates represent SD change in outcome variable per SD change in the urinary biomarker tested. Binary outcomes: atrial fibrillation (AF), coronary artery disease (CAD), heart failure (HF), hemorrhagic stroke (HS), ischemic stroke (IS), lipid medications (LIP) and type 2 diabetes (T2D). Continuous outcomes: systolic and diastolic blood pressure (SBP, and DBP; respectively), body fat percentage (BF), body mass index (BMI), waist-to-hip ratio (WHR). Models are adjusted for sex, age, center in UK, ethnicity, smoking, blood pressure (SBP and DBP), medication, lipids, T2D, obesity (BMI, WHR, body fat percentage) alcohol, Townsend index. P-values significant after Bonferroni correction ($p \leq 4.17E-03$) are bold. N, incident events in the UK Biobank.

Abbreviations: HR, hazard ratio; OR, odds ratio; CI, confidence interval; P, p-value.

**eTable 4.** Observational associations of urinary sodium, potassium and albumin with cardiovascular risk factors in UK Biobank using multivariable-adjusted logistic regression for main analyses.

| Sodium | Beta | lower | upper | P |
|---|---|---|---|---|
| **BF** | 0.0022 | 0.0022 | 0.0023 | **<2e-16** |
| **BMI** | 0.0038 | 0.0037 | 0.0039 | **<2e-16** |
| **WHR** | 0.0022 | 0.0021 | 0.0022 | **<2e-16** |
| **Potassium** | | | | |
| **BF** | 0.0021 | 0.0020 | 0.0021 | **<2e-16** |
| **BMI** | 0.0034 | 0.0033 | 0.0034 | **<2e-16** |
| **WHR** | 0.0015 | 0.0014 | 0.0015 | **<2e-16** |
| **Albumin** | | | | |
| **BF** | 0.0001 | 0.0001 | 0.0002 | **1.46E-09** |
| **BMI** | 0.0003 | 0.0002 | 0.0003 | **<2e-16** |
| **WHR** | 0.0003 | 0.0002 | 0.0003 | **<2e-16** |

Continuous outcomes: body fat percentage (BF), body mass index (BMI), waist-to-hip ratio (WHR). Models are adjusted for sex, age, center in UK, ethnicity, smoking, blood pressure, medication, lipids, type two diabetes, obesity (BMI, WHR, body fat percentage) alcohol, Townsend index. Significant p-values in bold.

Abbreviations: P, p-value.

**eTable 5.** Sex-stratified associations of the urinary sodium to potassium ratio (UNa/UK), sodium and potassium adjusted for creatinine (UNa/UCr and UK/UCr), and urinary albumin to creatinine ratio (UAlb/UCr) with cardiovascular outcomes in UK Biobank using multivariable-adjusted Cox proportional hazards models, and multivariable-adjusted linear and logistic regression for main analysis.

| **UNa/UK** | | | | | | | | | | | |
|---|---|---|---|---|---|---|---|---|---|---|---|
| **Females** | | | | | | **Males** | | | | | |
| **Outcome** | **N** | **HR/OR/Beta** | **Lower** | **Higher** | **P** | **N** | **HR/OR/Beta** | **Lower** | **Higher** | **P** | **P interaction** |
| **AF** | 3,329 | 0.917 | 0.886 | 0.949 | **6.22E-07** | 5,867 | 0.887 | 0.863 | 0.911 | **<2e-16** | 4.25E-02 |
| **CAD** | 1,925 | 0.992 | 0.948 | 1.037 | 7.15E-01 | 5,450 | 0.949 | 0.923 | 0.976 | **2.78E-04** | 7.59E-03 |
| **HF** | 971 | 0.979 | 0.92 | 1.042 | 5.13E-01 | 1,804 | 0.946 | 0.902 | 0.993 | 2.30E-02 | 1.90E-01 |
| **HS** | 487 | 1.02 | 0.931 | 1.117 | 6.75E-01 | 491 | 0.94 | 0.857 | 1.033 | 1.98E-01 | 9.09E-02 |
| **IS** | 720 | 0.963 | 0.894 | 1.037 | 3.18E-01 | 1,168 | 0.904 | 0.851 | 0.96 | **1.00E-03** | 1.70E-01 |
| **LIPIDS** | 28,711 | 0.945 | 0.931 | 0.959 | **3.25E-13** | 39,186 | 0.937 | 0.923 | 0.950 | **<2e-16** | 1.24E-06 |
| **T2D** | 8,375 | 0.999 | 0.973 | 1.025 | 9.37E-01 | 11,501 | 0.939 | 0.917 | 0.961 | **8.73E-08** | 1.01E-03 |
| **SBP** | | 0.131 | 0.127 | 0.135 | **<2e-16** | | 0.116 | 0.112 | 0.120 | **<2e-16** | 9.14E-01 |
| **DBP** | | 0.072 | 0.068 | 0.076 | **<2e-16** | | 0.081 | 0.077 | 0.086 | **<2e-16** | 7.79E-05 |
| **BF** | | 0.017 | 0.014 | 0.020 | **<2e-16** | | 0.038 | 0.035 | 0.041 | **<2e-16** | 2.00E-03 |
| **BMI** | | 0.043 | 0.039 | 0.047 | **<2e-16** | | 0.058 | 0.054 | 0.062 | **<2e-16** | 1.11E-01 |
| **WHR** | | 0.035 | 0.032 | 0.038 | **<2e-16** | | 0.056 | 0.053 | 0.059 | **<2e-16** | 2.01E-04 |
| **UNa/UCr** | | | | | | | | | | | |
| **Females** | | | | | | **Males** | | | | | |
| **Outcome** | **N** | **HR/OR/Beta** | **Lower** | **Higher** | **P** | **N** | **HR/OR/Beta** | **Lower** | **Higher** | **P** | **P interaction** |
| **AF** | 3,341 | 0.914 | 0.886 | 0.943 | **2.30E-08** | 5,878 | 0.898 | 0.875 | 0.923 | **5.88E-15** | 1.83E-01 |
| **CAD** | 1,929 | 0.941 | 0.903 | 0.981 | 5.00E-03 | 5,458 | 0.933 | 0.907 | 0.96 | **1.30E-06** | 2.59E-01 |

| | | | | | | | | | | | |
|---|---|---|---|---|---|---|---|---|---|---|---|
| **HF** | 972 | 0.954 | 0.901 | 1.01 | 1.06E-01 | 1,804 | 0.925 | 0.882 | 0.969 | **1.00E-03** | 2.04E-01 |
| **HS** | 487 | 1.014 | 0.93 | 1.106 | 7.45E-01 | 491 | 0.888 | 0.808 | 0.975 | 1.30E-02 | 1.59E-02 |
| **IS** | 719 | 0.895 | 0.835 | 0.959 | **2.00E-03** | 1,169 | 0.865 | 0.814 | 0.919 | **2.68E-06** | 4.87E-01 |
| **LIPIDS** | 28,711 | 0.958 | 0.945 | 0.972 | **3.39E-09** | 39,186 | 0.960 | 0.946 | 0.974 | **3.97E-08** | 8.61E-01 |
| **T2D** | 8,375 | 0.986 | 0.962 | 1.010 | 2.56E-01 | 11,501 | 0.991 | 0.969 | 1.014 | 4.59E-01 | 3.17E-01 |
| **SBP** | | 0.120 | 0.117 | 0.124 | **<2e-16** | | 0.119 | 0.115 | 0.123 | **<2e-16** | 6.89E-01 |
| **DBP** | | 0.073 | 0.069 | 0.076 | **<2e-16** | | 0.087 | 0.082 | 0.091 | **<2e-16** | 6.08E-05 |
| **BF** | | -0.050 | -0.053 | -0.047 | **<2e-16** | | -0.023 | -0.026 | -0.020 | **<2e-16** | <2e-16 |
| **BMI** | | -0.066 | -0.070 | -0.063 | **<2e-16** | | -0.034 | -0.037 | -0.030 | **<2e-16** | <2e-16 |
| **WHR** | | -0.012 | -0.015 | -0.009 | **5.21E-16** | | 0.002 | -0.001 | 0.005 | 2.52E-01 | 1.66E-05 |
| **UK/UCr** | | | | | | | | | | | |
| **Females** | | | | | | **Males** | | | | | |
| **Outcome** | **N** | **HR/OR/Beta** | **Lower** | **Higher** | **P** | **N** | **HR/OR/Beta** | **Lower** | **Higher** | **P** | **P interaction** |
| **AF** | 3,349 | 0.969 | 0.934 | 1.006 | 9.70E-02 | 5,880 | 0.998 | 0.967 | 1.029 | 8.79E-01 | 2.20E-01 |
| **CAD** | 1,932 | 0.912 | 0.869 | 0.958 | **2.23E-04** | 5,453 | 0.964 | 0.934 | 0.995 | 2.50E-02 | 2.09E-02 |
| **HF** | 978 | 0.913 | 0.852 | 0.978 | 9.00E-03 | 1,811 | 0.919 | 0.869 | 0.971 | **3.00E-03** | 8.76E-01 |
| **HS** | 488 | 0.983 | 0.894 | 1.081 | 7.18E-01 | 491 | 0.901 | 0.811 | 1 | 5.00E-02 | 3.30E-01 |
| **IS** | 725 | 0.867 | 0.801 | 0.939 | **4.14E-04** | 1,169 | 0.911 | 0.851 | 0.977 | 8.00E-03 | 2.76E-01 |
| **LIPIDS** | 28,711 | 1.018 | 1.001 | 1.035 | 3.71E-02 | 39,186 | 1.046 | 1.029 | 1.064 | **8.38E-08** | 3.19E-11 |
| **T2D** | 8,375 | 0.971 | 0.943 | 1.000 | 5.39E-02 | 11,501 | 1.103 | 1.073 | 1.134 | **2.06E-12** | 3.89E-08 |
| **SBP** | | -0.014 | -0.018 | -0.010 | **1.78E-12** | | -0.006 | -0.010 | -0.001 | 9.71E-03 | 3.28E-05 |
| **DBP** | | 0.001 | -0.003 | 0.005 | 6.32E-01 | | -0.0005 | -0.005 | 0.004 | 8.42E-01 | 1.57E-02 |
| **BF** | | -0.099 | -0.102 | -0.096 | **<2e-16** | | -0.095 | -0.099 | -0.092 | **<2e-16** | 6.76E-16 |
| **BMI** | | -0.160 | -0.164 | -0.156 | **<2e-16** | | -0.141 | -0.145 | -0.137 | **<2e-16** | <2e-16 |
| **WHR** | | -0.068 | -0.071 | -0.065 | **<2e-16** | | -0.086 | -0.090 | -0.083 | **<2e-16** | 1.50E-02 |

| **UAlb/UCr** | | | | | | | | | | | |
|---|---|---|---|---|---|---|---|---|---|---|---|
| **Females** | | | | | | **Males** | | | | | |
| **Outcome** | **N** | **HR/OR/Beta** | **Lower** | **Higher** | **P** | **N** | **HR/OR/Beta** | **Lower** | **Higher** | **P** | **P interaction** |
| **AF** | 3,362 | 1.125 | 1.08 | 1.172 | **1.28E-08** | 5,889 | 1.176 | 1.147 | 1.206 | **<2e-16** | 4.39E-01 |
| **CAD** | 1,938 | 1.066 | 1.011 | 1.123 | 1.80E-02 | 5,459 | 1.049 | 1.021 | 1.077 | **4.84E-04** | 1.90E-02 |
| **HF** | 980 | 1.243 | 1.156 | 1.336 | **4.40E-09** | 1,811 | 1.228 | 1.175 | 1.283 | **<2e-16** | 2.23E-01 |
| **HS** | 488 | 1.083 | 0.974 | 1.204 | 1.42E-01 | 491 | 1.201 | 1.1 | 1.312 | **4.61E-05** | 3.44E-01 |
| **IS** | 725 | 1.027 | 0.942 | 1.12 | 5.44E-01 | 1,170 | 1.077 | 1.017 | 1.141 | 1.10E-02 | 5.05E-01 |
| **LIPIDS** | 28,711 | 1.056 | 1.037 | 1.075 | **4.14E-09** | 39,186 | 1.065 | 1.050 | 1.080 | **<2e-16** | 7.93E-15 |
| **T2D** | 8,375 | 1.116 | 1.083 | 1.150 | **<2e-16** | 11,501 | 1.316 | 1.288 | 1.345 | **<2e-16** | <2e-16 |
| **SBP** | | 0.171 | 0.167 | 0.175 | **<2e-16** | | 0.150 | 0.147 | 0.154 | **<2e-16** | <2e-16 |
| **DBP** | | 0.139 | 0.134 | 0.143 | **<2e-16** | | 0.137 | 0.133 | 0.142 | **<2e-16** | 6.95E-04 |
| **BF** | | -0.152 | -0.156 | -0.149 | **<2e-16** | | -0.059 | -0.062 | -0.056 | **<2e-16** | <2e-16 |
| **BMI** | | -0.232 | -0.237 | -0.228 | **<2e-16** | | -0.099 | -0.103 | -0.095 | **<2e-16** | <2e-16 |
| **WHR** | | -0.087 | -0.091 | -0.084 | **<2e-16** | | -0.033 | -0.036 | -0.030 | **<2e-16** | <2e-16 |

Sex-stratified associations in the main cohort were performed excluding participants with kidney disease and/or cardiovascular disease at baseline. Estimates are from multivariable-adjusted linear (continuous risk factors), Cox proportional hazards models (cardiovascular outcomes) or logistic (lipid treatment and T2D) regression. The estimates represent SD change in outcome variable per SD change in urinary biomarkers. Binary outcomes: atrial fibrillation (AF), coronary artery disease (CAD), heart failure (HF), hemorrhagic stroke (HS), ischemic stroke (IS), lipid medications (LIP) and type 2 diabetes (T2D). Continuous outcomes: systolic and diastolic blood pressure (SBP, and DBP; respectively), body fat percentage (BF), body mass index (BMI), waist-to-hip ratio (WHR). Associations are adjusted for sex, age, center in UK, ethnicity, smoking, blood pressure [SBP and DBP], medication, lipids, T2D, obesity [BMI, WHR, body fat percentage] alcohol, Townsend index. P-values significant after Bonferroni correction ($p \leq 4.17E-03$) are bold. N, incident events in the UK Biobank.

Abbreviations: HR, hazard ratio; OR, odds ratio; CI, confidence interval; P, p-value.

**eTable 6.** Mendelian randomization analyses of associations of the urinary sodium to potassium ratio (UNa/UK), sodium and potassium adjusted for creatinine (UNa/UCr and UK/UCr) and urinary albumin to creatinine ratio (UAlb/UCr) with different cardiovascular risk factors and outcomes.

| **UNa/UK** | | | | | | | | |
|---|---|---|---|---|---|---|---|---|
| **Method** | **N SNP** | **Beta** | **SE** | **P** | **Global test P** | **Outlier corrected Beta** | **SD** | **P** |
| **SBP UKB** | | | | | <0.001 | n outliers = 8 | | |
| **IVW** | 26 | 2.118 | 1.876 | 2.59E-01 | | 2.630 | 1.093 | **2.59E-02** |
| **MR Egger** | 26 | -9.190 | 8.238 | 2.76E-01 | | | | |
| **WM** | 26 | 0.941 | 1.267 | 4.58E-01 | | | | |
| **SBP ICBP** | | | | | <0.001 | n outliers = 3 | | |
| **IVW** | 20 | 1.981 | 1.667 | 2.35E-01 | | 0.496 | 1.214 | 6.87E-01 |
| **MR Egger** | 20 | -4.085 | 6.521 | 5.39E-01 | | | | |
| **WM** | 20 | -0.044 | 1.541 | 9.77E-01 | | | | |
| **DBP UKB** | | | | | <0.001 | n outliers = 7 | | |
| **IVW** | 26 | 1.243 | 0.844 | 1.41E-01 | | 2.053 | 0.610 | **2.81E-03** |
| **MR Egger** | 26 | 1.121 | 3.856 | 7.74E-01 | | | | |
| **WM** | 26 | 0.828 | 0.626 | 1.86E-01 | | | | |
| **DBP ICBP** | | | | | 0.007 | n outliers = 1 | | |
| **IVW** | 20 | 1.194 | 0.832 | 1.51E-01 | | 0.591 | 0.713 | 4.16E-01 |
| **MR Egger** | 20 | 0.831 | 3.356 | 8.07E-01 | | | | |
| **WM** | 20 | 1.424 | 0.921 | 1.22E-01 | | | | |
| **BMI** | | | | | <0.001 | n outliers = 6 | | |
| **IVW** | 17 | 0.159 | 0.182 | 3.82E-01 | | 0.351 | 0.119 | **1.31E-02** |
| **MR Egger** | 17 | -0.033 | 1.056 | 9.76E-01 | | | | |

| | | | | | | | | |
|---|---|---|---|---|---|---|---|---|
| **WM** | 17 | 0.246 | 0.136 | 6.95E-02 | | | | |
| **WHR** | | | | | 0.039 | n outliers = 0 | | |
| **IVW** | 17 | 0.220 | 0.086 | 1.05E-02 | | | | |
| **MR Egger** | 17 | 0.736 | 0.480 | 1.46E-01 | | | | |
| **WM** | 17 | 0.240 | 0.094 | 1.10E-02 | | | | |
| **UNa/UCr** | | | | | | | | |
| **Method** | **N SNP** | **Beta** | **SE** | **P** | **Global test P** | **Outlier corrected Beta** | **SD** | **P** |
| **SBP UKB** | | | | | <0.001 | n outliers = 7 | | |
| **IVW** | 24 | 3.233 | 1.768 | 6.75E-02 | | 3.998 | 1.067 | **1.61E-03** |
| **MR Egger** | 24 | 2.375 | 6.432 | 7.16E-01 | | | | |
| **WM** | 24 | 3.265 | 1.159 | 4.85E-03 | | | | |
| **SBP ICBP** | | | | | <0.001 | n outliers = 3 | | |
| **IVW** | 20 | 2.660 | 1.551 | 8.62E-02 | | 2.990 | 1.160 | **1.90E-02** |
| **MR Egger** | 20 | 3.252 | 5.509 | 5.62E-01 | | | | |
| **WM** | 20 | 4.862 | 1.395 | 4.90E-04 | | | | |
| **DBP UKB** | | | | | <0.001 | n outliers = 4 | | |
| **IVW** | 24 | 1.663 | 0.716 | 2.01E-02 | | 2.295 | 0.590 | **9.12E-04** |
| **MR Egger** | 24 | 3.794 | 2.561 | 1.53E-01 | | | | |
| **WM** | 24 | 0.818 | 0.644 | 2.05E-01 | | | | |
| **DBP ICBP** | | | | | 0.003 | n outliers = 1 | | |
| **IVW** | 20 | 0.956 | 0.769 | 2.14E-01 | | 0.657 | 0.694 | 3.56E-01 |
| **MR Egger** | 20 | 3.472 | 2.670 | 2.10E-01 | | | | |
| **WM** | 20 | 2.232 | 0.837 | 7.68E-03 | | | | |
| **BMI** | | | | | <0.001 | n outliers = 1 | | |
| **IVW** | 14 | -0.289 | 0.098 | 3.20E-03 | | -0.368 | 0.072 | **2.60E-04** |

| | | | | | | | | |
|---|---|---|---|---|---|---|---|---|
| **MR Egger** | 14 | -0.286 | 0.432 | 5.20E-01 | | | | |
| **WM** | 14 | -0.389 | 0.081 | 1.60E-06 | | | | |
| **WHR** | | | | | 0.542 | n outliers = 0 | | |
| **IVW** | 14 | 0.235 | 0.061 | 1.08E-04 | | | | |
| **MR Egger** | 14 | 0.250 | 0.257 | 3.50E-01 | | | | |
| **WM** | 14 | 0.217 | 0.086 | 1.14E-02 | | | | |
| **UK/UCr** | | | | | | | | |
| **Method** | **N SNP** | **Beta** | **SE** | **P** | **Global test P** | **Outlier corrected Beta** | **SD** | **P** |
| **SBP UKB** | | | | | <0.001 | n outliers = 2 | | |
| **IVW** | 20 | 1.998 | 1.221 | 1.02E-01 | | 0.694 | 1.140 | 5.50E-01 |
| **MR Egger** | 20 | 5.694 | 4.381 | 2.10E-01 | | | | |
| **WM** | 20 | 1.790 | 1.163 | 1.24E-01 | | | | |
| **SBP ICBP** | | | | | <0.001 | n outliers = 2 | | |
| **IVW** | 17 | 2.195 | 1.998 | 2.72E-01 | | 2.941 | 1.552 | 7.89E-02 |
| **MR Egger** | 17 | -3.643 | 7.707 | 6.43E-01 | | | | |
| **WM** | 17 | 5.035 | 1.613 | 1.80E-03 | | | | |
| **DBP UKB** | | | | | <0.001 | n outliers = 1 | | |
| **IVW** | 20 | 0.456 | 0.604 | 4.51E-01 | | -0.097 | 0.554 | 8.63E-01 |
| **MR Egger** | 20 | -0.171 | 2.209 | 9.39E-01 | | | | |
| **WM** | 20 | 0.532 | 0.618 | 3.89E-01 | | | | |
| **DBP ICBP** | | | | | <0.001 | n outliers = 3 | | |
| **IVW** | 17 | 0.700 | 1.357 | 6.06E-01 | | 2.091 | 0.974 | 5.12E-02 |
| **MR Egger** | 17 | -5.809 | 5.054 | 2.68E-01 | | | | |
| **WM** | 17 | 2.127 | 0.985 | 3.08E-02 | | | | |
| **BMI** | | | | | <0.001 | n outliers = 3 | | |

| | | | | | | | | |
|---|---|---|---|---|---|---|---|---|
| **IVW** | 11 | -0.300 | 0.187 | 1.09E-01 | | -0.338 | 0.147 | 6.17E-02 |
| **MR Egger** | 11 | -0.906 | 0.726 | 2.44E-01 | | | | |
| **WM** | 11 | -0.115 | 0.118 | 3.30E-01 | | | | |
| **WHR** | | | | | | n outliers = 1 | | |
| **IVW** | 11 | 0.072 | 0.100 | 4.68E-01 | 0.028 | 0.027 | 0.080 | 7.45E-01 |
| **MR Egger** | 11 | -0.054 | 0.410 | 8.98E-01 | | | | |
| **WM** | 11 | 0.063 | 0.099 | 5.29E-01 | | | | |
| **UAlb/UCr** | | | | | | | | |
| **Method** | **N SNP** | **Beta** | **SE** | **P** | **Global test P** | **Outlier corrected Beta** | **SD** | **P** |
| **SBP UKB** | | | | | <0.001 | n outliers = 5 | | |
| **IVW** | 42 | 3.355 | 0.927 | 2.95E-04 | | 2.753 | 0.671 | **2.06E-04** |
| **MR Egger** | 42 | 1.126 | 2.289 | 6.26E-01 | | | | |
| **WM** | 42 | 2.174 | 0.757 | 4.07E-03 | | | | |
| **SBP ICBP** | | | | | <0.001 | n outliers = 4 | | |
| **IVW** | 36 | 2.186 | 1.311 | 9.53E-02 | | 2.296 | 1.026 | **3.18E-02** |
| **MR Egger** | 36 | -0.407 | 3.575 | 9.10E-01 | | | | |
| **WM** | 36 | 1.372 | 1.151 | 2.34E-01 | | | | |
| **DBP UKB** | | | | | <0.001 | n outliers = 7 | | |
| **IVW** | 42 | 0.974 | 0.553 | 7.83E-02 | | 0.848 | 0.393 | **3.77E-02** |
| **MR Egger** | 42 | 0.984 | 1.386 | 4.82E-01 | | | | |
| **WM** | 42 | 0.967 | 0.455 | 3.34E-02 | | | | |
| **DBP ICBP** | | | | | <0.001 | n outliers = 6 | | |
| **IVW** | 36 | 0.602 | 0.854 | 4.80E-01 | | 0.809 | 0.645 | 2.18E-01 |
| **MR Egger** | 36 | -1.275 | 2.329 | 5.88E-01 | | | | |
| **WM** | 36 | 0.260 | 0.695 | 7.08E-01 | | | | |

| | | | | | | | | |
|---|---|---|---|---|---|---|---|---|
| **BMI** | | | | | <0.001 | n outliers = 2 | | |
| **IVW** | 29 | -0.134 | 0.071 | 5.90E-02 | | -0.056 | 0.065 | 4.02E-01 |
| **MR Egger** | 29 | 0.103 | 0.346 | 7.68E-01 | | | | |
| **WM** | 29 | -0.025 | 0.075 | 7.44E-01 | | | | |
| **WHR** | | | | | 0.178 | n outliers = 0 | | |
| **IVW** | 29 | 0.187 | 0.055 | 7.36E-04 | | | | |
| **MR Egger** | 29 | 0.359 | 0.266 | 1.90E-01 | | | | |
| **WM** | 29 | 0.215 | 0.076 | 4.83E-03 | | | | |
| **T2D** | | | | | 0.003 | n outliers = 1 | | |
| **IVW** | 26 | 0.411 | 0.142 | 3.76E-03 | | 0.282 | 0.117 | **2.30E-02** |
| **MR Egger** | 26 | 0.432 | 0.730 | 5.60E-01 | | | | |
| **WM** | 26 | 0.208 | 0.160 | 1.94E-01 | | | | |
| **AF** | | | | | <0.001 | n outliers = 4 | | |
| **IVW** | 41 | 0.095 | 0.120 | 4.28E-01 | | 0.105 | 0.086 | 2.32E-01 |
| **MR Egger** | 41 | 0.225 | 0.317 | 4.83E-01 | | | | |
| **WM** | 41 | 0.058 | 0.113 | 6.09E-01 | | | | |

Three separate methods were used to estimate causal effects of urinary biomarkers with different risk factors for cardiovascular disease: the standard inverse-variance weighted (IVW) regression; and two robust regression methods, the weighted median-based method (WM), and Egger regression. We applied the global test to correct for pleiotropy. Significant p-values ($p \leq 0.05$) in bold.

Abbreviations: SBP, systolic blood pressure; DBP, diastolic blood pressure; BMI, body mass index; WHR, waist-to-hip ratio; T2D, type two diabetes; AF, atrial fibrillation; N, number of SNPs; SE, standard error; SD, standard deviation; P, P-value.

**eTable 7.** Mendelian randomization analyses of associations of systolic and diastolic blood pressure in the UK Biobank (SBP and DBP, respectively) and type two diabetes (T2D) with urinary sodium to potassium ratio (UNa/UK) and urinary albumin to creatinine ratio (UAlb/UCr).

| **SBP** | | | | | | | | |
|---|---|---|---|---|---|---|---|---|
| **Method** | **N SNP** | **Beta** | **SE** | **P** | **Global test P** | **Outlier corrected Beta** | **SE** | **P** |
| **UNa/UK** | | | | | <0.001 | n outliers = 11 | | |
| **IVW** | 290 | 0.0015 | 0.0007 | 3.42E-02 | | 0.0018 | 0.0006 | **2.68E-03** |
| **MR Egger** | 290 | -0.0089 | 0.0023 | 1.44E-04 | | | | |
| **WM** | 290 | 0.0008 | 0.0008 | 2.75E-01 | | | | |
| **UAlb/UCr** | | | | | <0.001 | n outliers = 12 | | |
| **IVW** | 290 | 0.0067 | 0.0007 | 2.00E-21 | | 0.0067 | 0.0006 | **9.48E-24** |
| **MR Egger** | 290 | 0.0063 | 0.0023 | 7.14E-03 | | | | |
| **WM** | 290 | 0.0069 | 0.0008 | 1.26E-19 | | | | |
| **DBP** | | | | | | | | |
| **Method** | **N SNP** | **Beta** | **SE** | **P** | **Global test P** | **Outlier corrected Beta** | **SE** | **P** |
| **UNa/UK** | | | | | <0.001 | n outliers = 12 | | |
| **IVW** | 253 | -0.0013 | 0.0013 | 3.15E-01 | | -0.0006 | 0.0011 | 5.77E-01 |
| **MR Egger** | 253 | -0.0140 | 0.0043 | 1.41E-03 | | | | |
| **WM** | 253 | 0.0003 | 0.0014 | 8.55E-01 | | | | |
| **UAlb/UCr** | | | | | <0.001 | n outliers = 12 | | |
| **IVW** | 253 | 0.0084 | 0.0014 | 8.08E-10 | | 0.0083 | 0.0012 | **9.23E-12** |
| **MR Egger** | 253 | 0.0083 | 0.0048 | 8.48E-02 | | | | |
| **WM** | 253 | 0.0090 | 0.0014 | 3.37E-11 | | | | |
| **T2D** | | | | | | | | |

| Method | N SNP | Beta | SE | P | Global test P | Outlier corrected Beta | SE | P |
|---|---|---|---|---|---|---|---|---|
| **UAlb/UCr** | | | | | <0.001 | n outliers = 5 | | |
| **IVW** | 156 | 0.0193 | 0.0051 | 1.42E-04 | | 0.0210 | 0.0042 | **1.89E-06** |
| **MR Egger** | 156 | 0.0239 | 0.0126 | 5.98E-02 | | | | |
| **WM** | 156 | 0.0253 | 0.0064 | 7.04E-05 | | | | |

Three separate methods were used to estimate causal effects of urinary biomarkers with different risk factors for cardiovascular disease: the standard inverse-variance weighted (IVW) regression; and two robust regression methods, the weighted median-based method (WM), and Egger regression. We applied the global test to correct for pleiotropy. Significant p-values ($p \leq 0.05$) in bold.

Abbreviations: N, number of SNPs; SE, standard error; SD, standard deviation; P, P-value.

**eTable 8.** Association of urinary albumin to creatinine ratio (UAlb/UCr), systolic blood pressure (SBP) and waist-to-hip ratio (WHR) with type 2 diabetes (T2D) and atrial fibrillation (AF) from mediation MR analyses to address whether UAlb/UCr had a causal effect on T2D and AF, independently of SBP and WHR.

| **T2D** | **Beta** | **SE** | **P** |
|---|---|---|---|
| **UAlb/UCr** | 0.4809 | 0.1987 | **1.55E-02** |
| **SBP** | 0.0195 | 0.0044 | **9.06E-06** |
| **WHR** | 0.1339 | 0.1018 | 1.88E-01 |
| **AF** | | | |
| **UAlb/UCr** | 0.0951 | 0.1521 | 5.32E-01 |
| **SBP** | 0.0124 | 0.0035 | **3.19E-04** |
| **WHR** | -0.1970 | 0.0825 | **1.70E-02** |

Estimates are from multivariable MR regression-based model. Significant p-values ($p \leq 0.05$) in bold.

Abbreviations: SE, standard error; P, P-Value.

### Supplemental Figures

**eFigure 1.** Association results (BETA/HR/OR) of urinary sodium to potassium ratio (Na/UK), urinary sodium to creatinine ratio (UNa/UCr), urinary potassium to creatinine ratio (UK/UCr) and urinary albumin to creatinine ratio (UAlb/UCr) with cardiovascular outcomes in UK Biobank using multivariable-adjusted linear and logistic regression, and multivariable-adjusted Cox proportional hazards models. a. Binary outcomes: atrial fibrillation (AF); coronary artery disease (CAD); heart failure (HF); hemorrhagic and ischemic stroke (HS and IS, respectively); lipid medications (LIP) and type 2 diabetes (T2D). b. Continuous outcomes: systolic and diastolic blood pressure (SBP and DBP, respectively), body fat percentage (BF); body mass index (BMI), waist-to-hip ratio (WHR).

a.

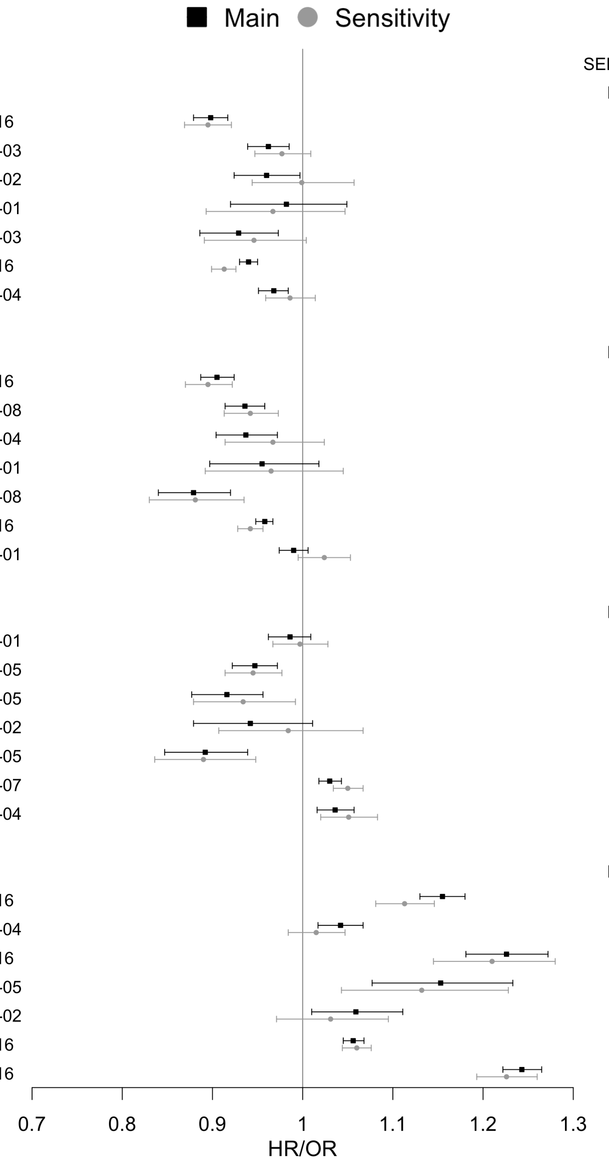

| | | MAIN Events | MAIN HR/OR | MAIN P | SENSITIVITY Events | SENSITIVITY HR/OR | SENSITIVITY P |
|---|---|---|---|---|---|---|---|
| UNa/UK | AF | 9196 | 0.898 | <2e-16 | 5499 | 0.895 | 5.03E-14 |
| | CAD | 7375 | 0.962 | 1.24E-03 | 4659 | 0.977 | 1.53E-01 |
| | HF | 2775 | 0.96 | 3.41E-02 | 1422 | 0.999 | 9.72E-01 |
| | HS | 978 | 0.982 | 5.91E-01 | 714 | 0.967 | 4.07E-01 |
| | IS | 1888 | 0.929 | 1.94E-03 | 1268 | 0.946 | 6.69E-02 |
| | LIPID | 67898 | 0.94 | <2e-16 | 28978 | 0.913 | <2e-16 |
| | T2D | 19876 | 0.968 | 1.84E-04 | 7758 | 0.986 | 3.16E-01 |
| | | Events | HR/OR | P | Events | HR/OR | P |
| UNa/UCr | AF | 9219 | 0.905 | <2e-16 | 5516 | 0.895 | 8.48E-14 |
| | CAD | 7387 | 0.936 | 1.93E-08 | 4665 | 0.942 | 2.37E-04 |
| | HF | 2776 | 0.937 | 4.76E-04 | 1423 | 0.967 | 2.47E-01 |
| | HS | 978 | 0.955 | 1.60E-01 | 714 | 0.965 | 3.82E-01 |
| | IS | 1888 | 0.879 | 3.04E-08 | 1268 | 0.881 | 3.26E-05 |
| | LIPID | 67898 | 0.958 | <2e-16 | 28978 | 0.942 | 2.46E-15 |
| | T2D | 19876 | 0.99 | 2.25E-01 | 7758 | 1.024 | 1.02E-01 |
| | | Events | HR/OR | P | Events | HR/OR | P |
| UK/UCr | AF | 9229 | 0.986 | 2.35E-01 | 5513 | 0.997 | 8.50E-01 |
| | CAD | 7385 | 0.947 | 5.46E-05 | 4666 | 0.945 | 9.22E-04 |
| | HF | 2789 | 0.916 | 6.07E-05 | 1425 | 0.934 | 2.52E-02 |
| | HS | 979 | 0.942 | 9.83E-02 | 715 | 0.984 | 6.95E-01 |
| | IS | 1894 | 0.892 | 1.57E-05 | 1270 | 0.89 | 2.89E-04 |
| | LIPID | 67898 | 1.03 | 6.52E-07 | 28978 | 1.05 | 8.47E-10 |
| | T2D | 19876 | 1.036 | 4.82E-04 | 7758 | 1.051 | 1.10E-03 |
| | | Events | HR/OR | P | Events | HR/OR | P |
| UAlb/UCr | AF | 9251 | 1.155 | <2e-16 | 5529 | 1.113 | 4.40E-13 |
| | CAD | 7397 | 1.042 | 6.87E-04 | 4672 | 1.015 | 3.56E-01 |
| | HF | 2791 | 1.226 | <2e-16 | 1425 | 1.21 | 1.61E-11 |
| | HS | 979 | 1.153 | 3.90E-05 | 715 | 1.132 | 2.95E-03 |
| | IS | 1895 | 1.059 | 1.82E-02 | 1271 | 1.031 | 3.22E-01 |
| | LIPID | 67898 | 1.056 | <2e-16 | 28978 | 1.06 | 2.61E-14 |
| | T2D | 19876 | 1.243 | <2e-16 | 7758 | 1.226 | <2e-16 |

b.

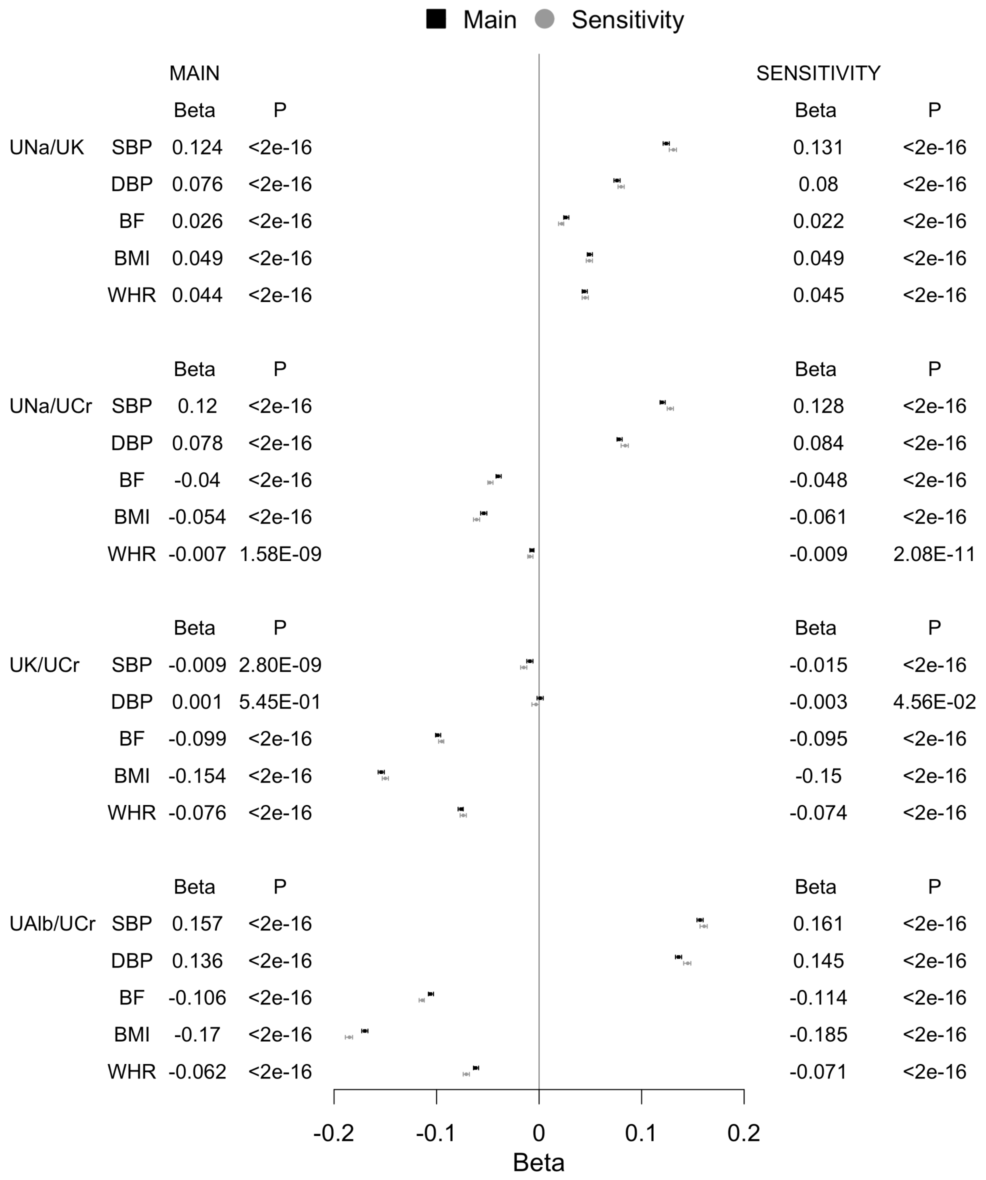

| | | MAIN Beta | MAIN P | SENSITIVITY Beta | SENSITIVITY P |
|---|---|---|---|---|---|
| UNa/UK | SBP | 0.124 | <2e-16 | 0.131 | <2e-16 |
| | DBP | 0.076 | <2e-16 | 0.08 | <2e-16 |
| | BF | 0.026 | <2e-16 | 0.022 | <2e-16 |
| | BMI | 0.049 | <2e-16 | 0.049 | <2e-16 |
| | WHR | 0.044 | <2e-16 | 0.045 | <2e-16 |
| UNa/UCr | SBP | 0.12 | <2e-16 | 0.128 | <2e-16 |
| | DBP | 0.078 | <2e-16 | 0.084 | <2e-16 |
| | BF | -0.04 | <2e-16 | -0.048 | <2e-16 |
| | BMI | -0.054 | <2e-16 | -0.061 | <2e-16 |
| | WHR | -0.007 | 1.58E-09 | -0.009 | 2.08E-11 |
| UK/UCr | SBP | -0.009 | 2.80E-09 | -0.015 | <2e-16 |
| | DBP | 0.001 | 5.45E-01 | -0.003 | 4.56E-02 |
| | BF | -0.099 | <2e-16 | -0.095 | <2e-16 |
| | BMI | -0.154 | <2e-16 | -0.15 | <2e-16 |
| | WHR | -0.076 | <2e-16 | -0.074 | <2e-16 |
| UAlb/UCr | SBP | 0.157 | <2e-16 | 0.161 | <2e-16 |
| | DBP | 0.136 | <2e-16 | 0.145 | <2e-16 |
| | BF | -0.106 | <2e-16 | -0.114 | <2e-16 |
| | BMI | -0.17 | <2e-16 | -0.185 | <2e-16 |
| | WHR | -0.062 | <2e-16 | -0.071 | <2e-16 |

Main analyses were performed excluding participants with kidney disease and/or cardiovascular disease at baseline. Sensitivity analyses was performed excluding participants with kidney disease and/or cardiovascular disease at baseline and/or participants using medications affecting renal excretion. The betas from linear regression represent SD change in outcome variable per SD change in urinary biomarkers. Model adjustment: sex, age, region of the UK Biobank assessment center, ethnicity, smoking, alcohol, physical activity, Townsend index, blood pressure (DBP and SBP), obesity (BMI, body fat percentage, WHR), lipid medications, T2D, and medications affecting renal excretion. Abbreviations: HR, hazard ratio; OR, odds ratio.

**eFigure 2.** Relations of the urinary sodium to potassium ratio (UNa/UK) with a. atrial fibrillation (AF); b. coronary artery disease (CAD); c. heart failure (HF); d. hemorrhagic stroke (HS); e. ischemic stroke (IS). Lines are based on a regression spline of Cox proportional hazards.

a.

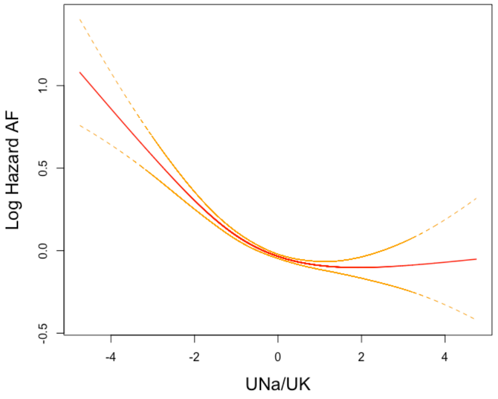

b.

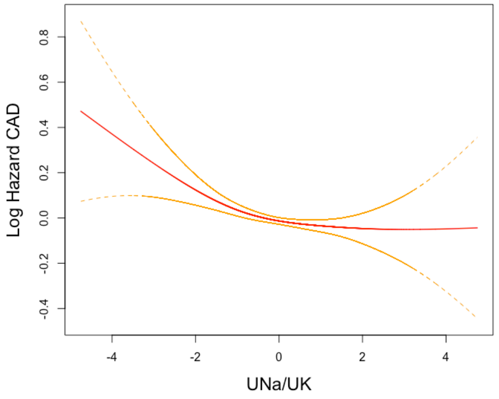

c .

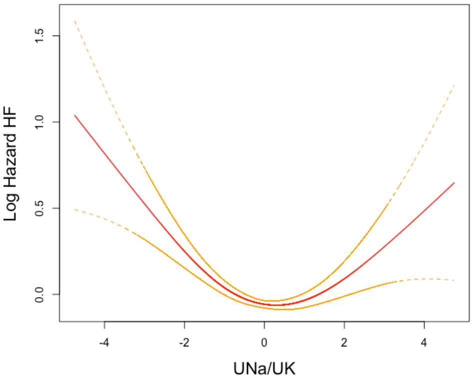

d.

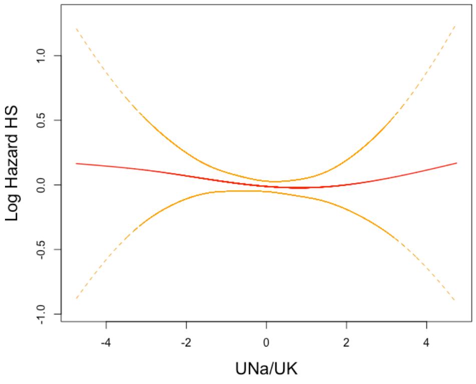

e.

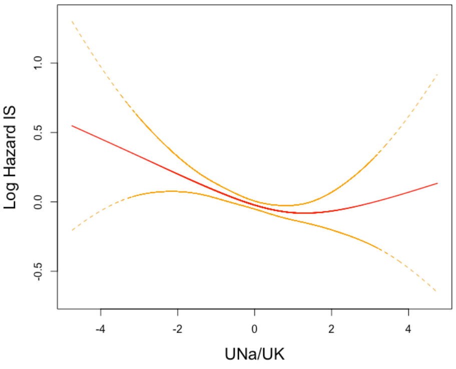

**eFigure 3.** Relations of the urinary albumin to creatinine ratio (UAlb/UCr) with a. atrial fibrillation (AF); b. coronary artery disease (CAD); c. heart failure (HF); d. hemorrhagic stroke (HS); e. ischemic stroke (IS). Lines are based on a regression spline of Cox proportional hazards.

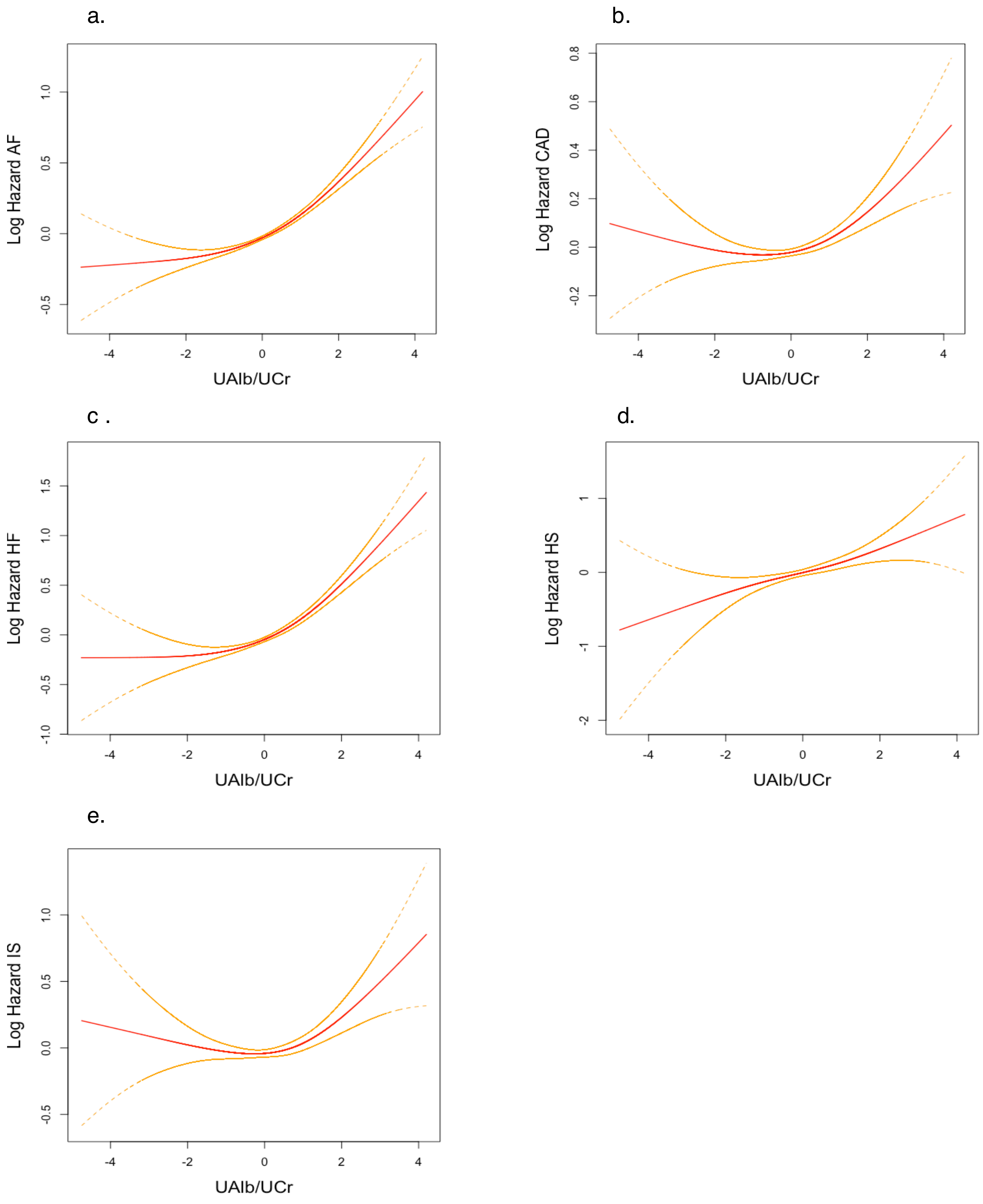

**eFigure 4.** Association results (BETA/HR/OR) of urinary sodium to potassium ratio (Na/UK), urinary sodium to creatinine ratio (UNa/UCr), urinary potassium to creatinine ratio (UK/UCr) and urinary albumin to creatinine ratio (UAlb/UCr) stratified by sex with cardiovascular outcomes in UK Biobank using multivariable-adjusted linear and logistic regression, and multivariable-adjusted Cox proportional hazards models in main analyses. a. Binary outcomes: atrial fibrillation (AF); coronary artery disease (CAD); heart failure (HF); hemorrhagic and ischemic stroke (HS and IS, respectively); lipid medications (LIP) and type 2 diabetes (T2D). b. Continuous outcomes: systolic and diastolic blood pressure (SBP and DBP, respectively), body fat percentage (BF); body mass index (BMI), waist-to-hip ratio (WHR).

a.

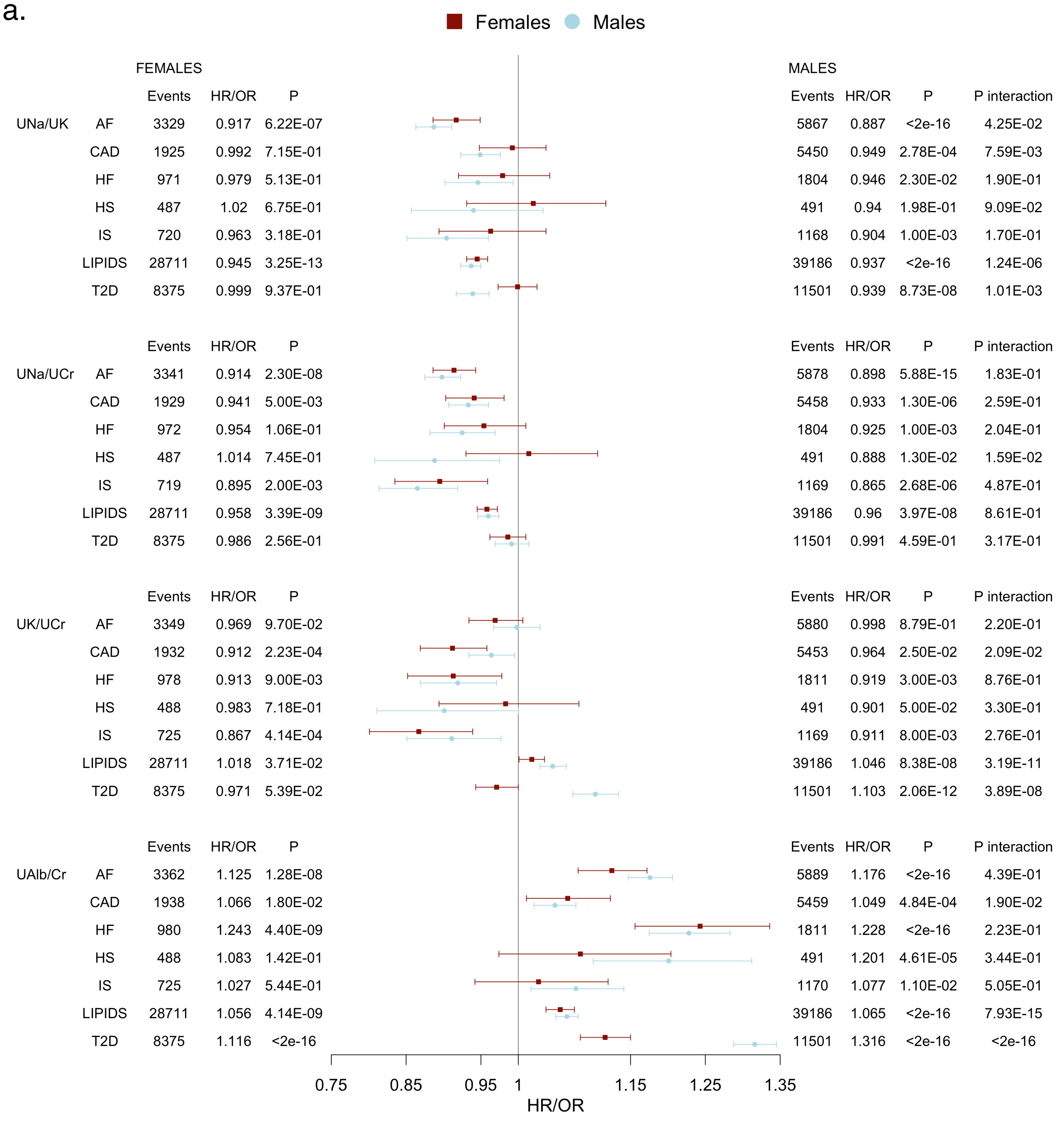

b.

| | | FEMALES Beta | FEMALES P | MALES Beta | MALES P | P interaction |
|---|---|---|---|---|---|---|
| UNa/UK | SBP | 0.131 | <2e-16 | 0.116 | <2e-16 | 9.14E-01 |
| | DBP | 0.072 | <2e-16 | 0.081 | <2e-16 | 7.79E-05 |
| | BF | 0.017 | <2e-16 | 0.038 | <2e-16 | 2.00E-03 |
| | BMI | 0.043 | <2e-16 | 0.058 | <2e-16 | 1.11E-01 |
| | WHR | 0.035 | <2e-16 | 0.056 | <2e-16 | 2.01E-04 |
| UNa/UCr | SBP | 0.12 | <2e-16 | 0.119 | <2e-16 | 6.89E-01 |
| | DBP | 0.073 | <2e-16 | 0.087 | 2.04e-315 | 6.08E-05 |
| | BF | -0.05 | <2e-16 | -0.023 | 1.33E-45 | <2e-16 |
| | BMI | -0.066 | <2e-16 | -0.034 | 3.29E-63 | <2e-16 |
| | WHR | -0.012 | 5.21E-16 | 0.002 | 2.52E-01 | 1.66E-05 |
| UK/UCr | SBP | -0.014 | 1.78E-12 | -0.006 | 9.71E-03 | 3.28E-05 |
| | DBP | 0.001 | 6.32E-01 | -0.0005 | 8.42E-01 | 1.57E-02 |
| | BF | -0.099 | <2e-16 | -0.095 | <2e-16 | 6.76E-16 |
| | BMI | -0.16 | <2e-16 | -0.141 | <2e-16 | <2e-16 |
| | WHR | -0.068 | <2e-16 | -0.086 | <2e-16 | 1.50E-02 |
| UAlb/Cr | SBP | 0.171 | <2e-16 | 0.15 | <2e-16 | <2e-16 |
| | DBP | 0.139 | <2e-16 | 0.137 | <2e-16 | 6.95E-04 |
| | BF | -0.152 | <2e-16 | -0.059 | <2e-16 | <2e-16 |
| | BMI | -0.232 | <2e-16 | -0.099 | <2e-16 | <2e-16 |
| | WHR | -0.087 | <2e-16 | -0.033 | <2e-16 | <2e-16 |

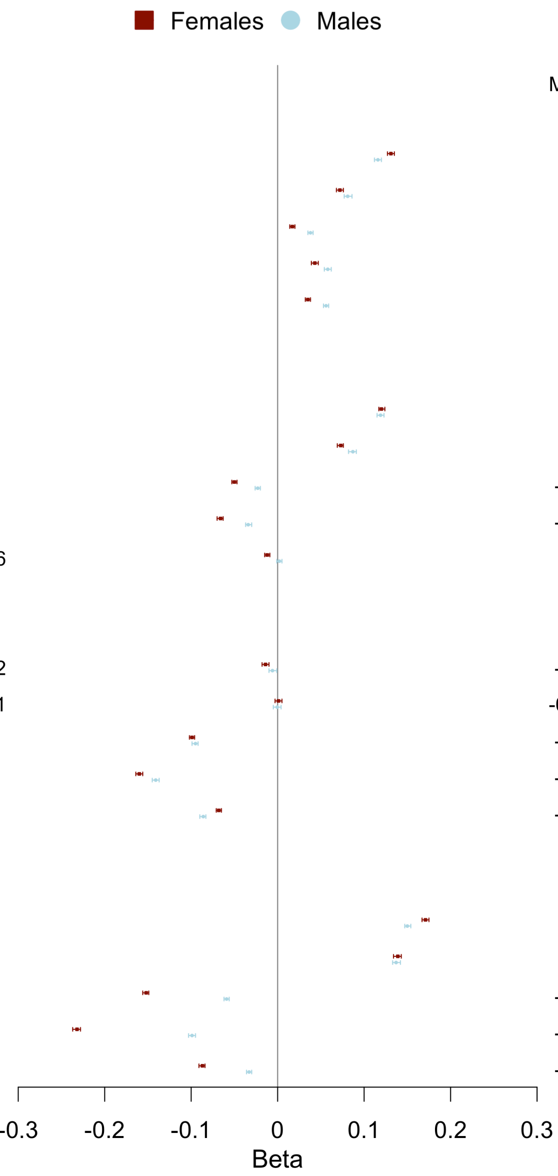

Sex-stratified analysis (N males= 212,534; N females= 265,777) performed in the main cohort excluding participants with diagnosis indicating decreased kidney function (N= 7,221) and/or cardiovascular disease at baseline (N= 17,087). The betas from linear regression represent SD change in outcome variable per SD change in UNa/UK. Model adjustment: sex, age, region of the UK Biobank assessment center, ethnicity, smoking, alcohol, physical activity, Townsend index, blood pressure (DBP and SBP), obesity (BMI, body fat percentage, WHR), lipid medications, T2D, and medications affecting renal excretion. Abbreviations: HR, hazard ratio; OR, odds ratio.

**eFigure 5.** Effects of genetic variants on urinary sodium to potassium ratio (UNa/UK) and risk of a. systolic blood pressure in UK Biobank (SBP UKB); b. systolic blood pressure from International Consortium of Blood Pressure (SBP ICBP); c. diastolic blood pressure in UK Biobank (DBP UKB); d. diastolic blood pressure from International Consortium of Blood Pressure (DBP ICBP); e. body mass index (BMI); f. waist-to-hip ratio (WHR). Variants showing significant heterogeneity are highlighted in red.

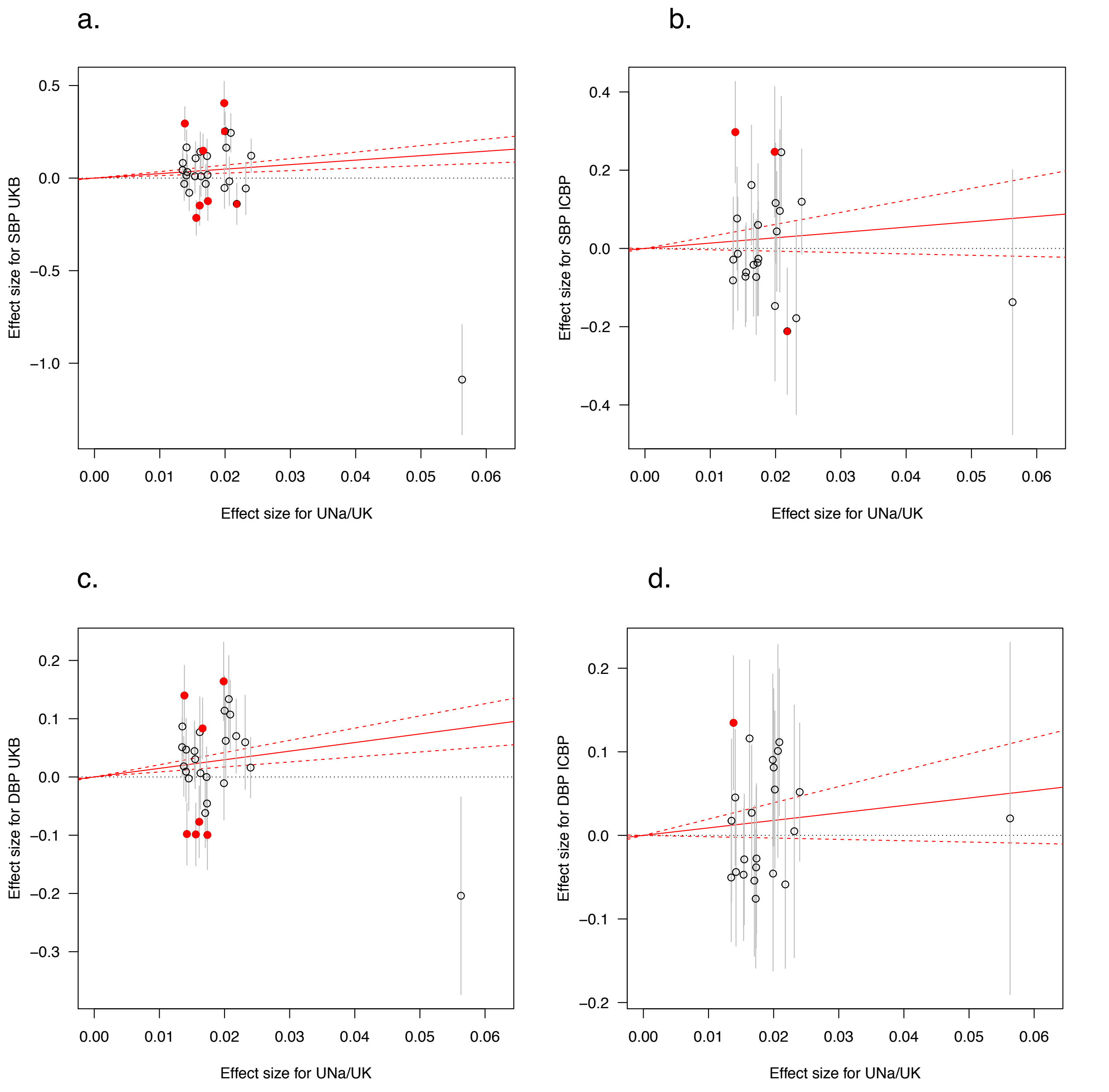

e.

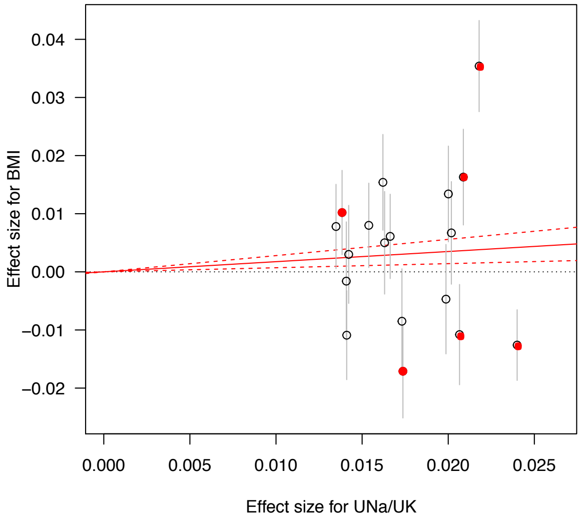
0.04
0.03
0.02
0.01
0.00
−0.01
−0.02
Effect size for BMI
0.000
0.005
0.010
0.015
0.020
0.025
Effect size for UNa/UK

f.

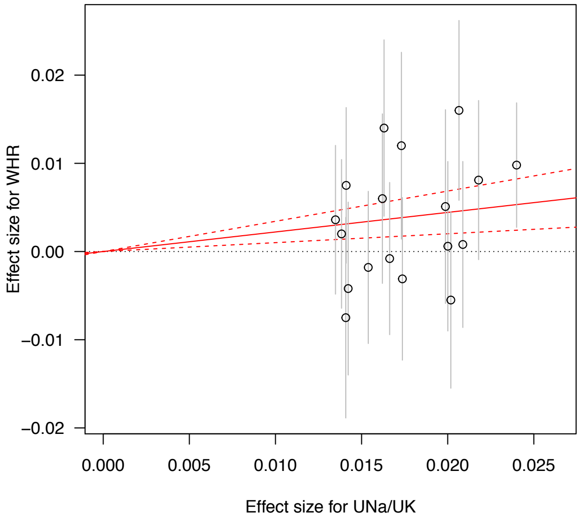
0.02
0.01
0.00
−0.01
−0.02
Effect size for WHR
0.000
0.005
0.010
0.015
0.020
0.025
Effect size for UNa/UK

**eFigure 6.** Effects of genetic variants on urinary sodium to creatinine ratio (UNa/UCr) and risk of a. systolic blood pressure in UK Biobank (SBP UKB); b. systolic blood pressure from International Consortium of Blood Pressure (SBP ICBP); c. diastolic blood pressure in UK Biobank (DBP UKB); d. diastolic blood pressure from International Consortium of Blood Pressure (DBP ICBP); e. body mass index (BMI); f. waist-to-hip ratio (WHR). Variants showing significant heterogeneity are highlighted in red.

a.

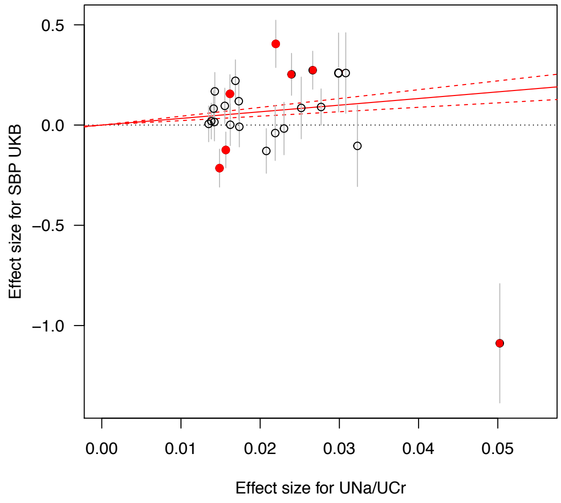

b.

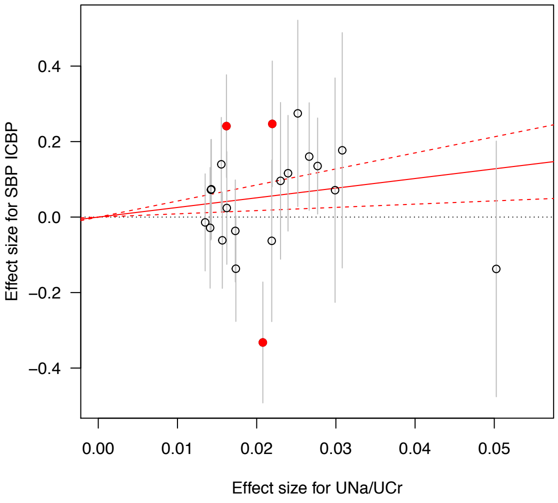

c.

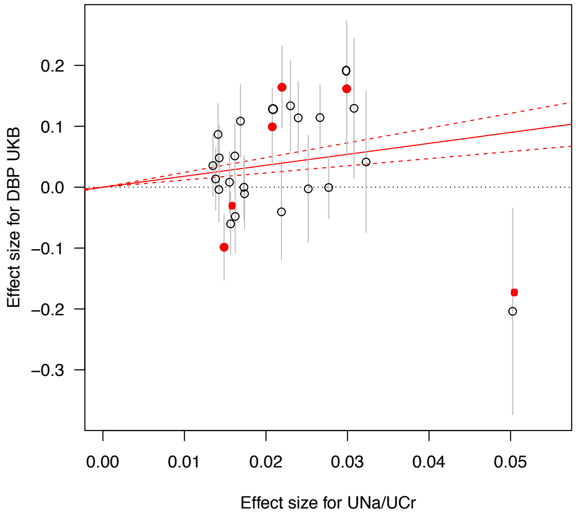

d.

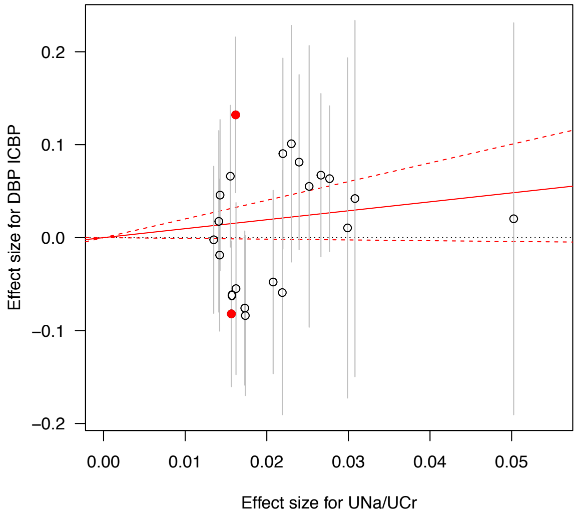

e.

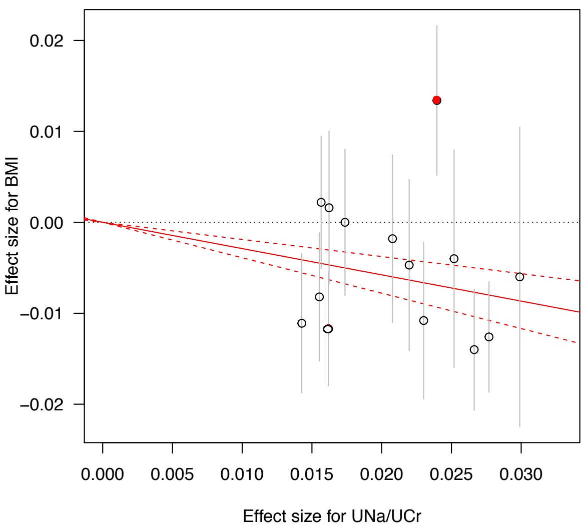
0.02
0.01
0.00
−0.01
−0.02
Effect size for BMI
0.000
0.005
0.010
0.015
0.020
0.025
0.030
Effect size for UNa/UCr

f.

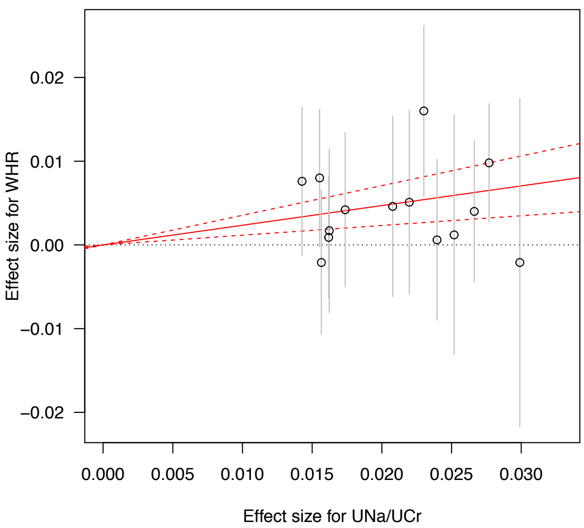
0.02
0.01
0.00
−0.01
−0.02
Effect size for WHR
0.000
0.005
0.010
0.015
0.020
0.025
0.030
Effect size for UNa/UCr

**eFigure 7.** Effects of genetic variants on urinary potassium to creatinine ratio (UK/UCr) and risk of a. systolic blood pressure in UK Biobank (SBP UKB); b. systolic blood pressure from International Consortium of Blood Pressure (SBP ICBP); c. diastolic blood pressure in UK Biobank (DBP UKB); d. diastolic blood pressure from International Consortium of Blood Pressure (DBP ICBP); e. body mass index (BMI); f. waist-to-hip ratio (WHR). Variants showing significant heterogeneity are highlighted in red.

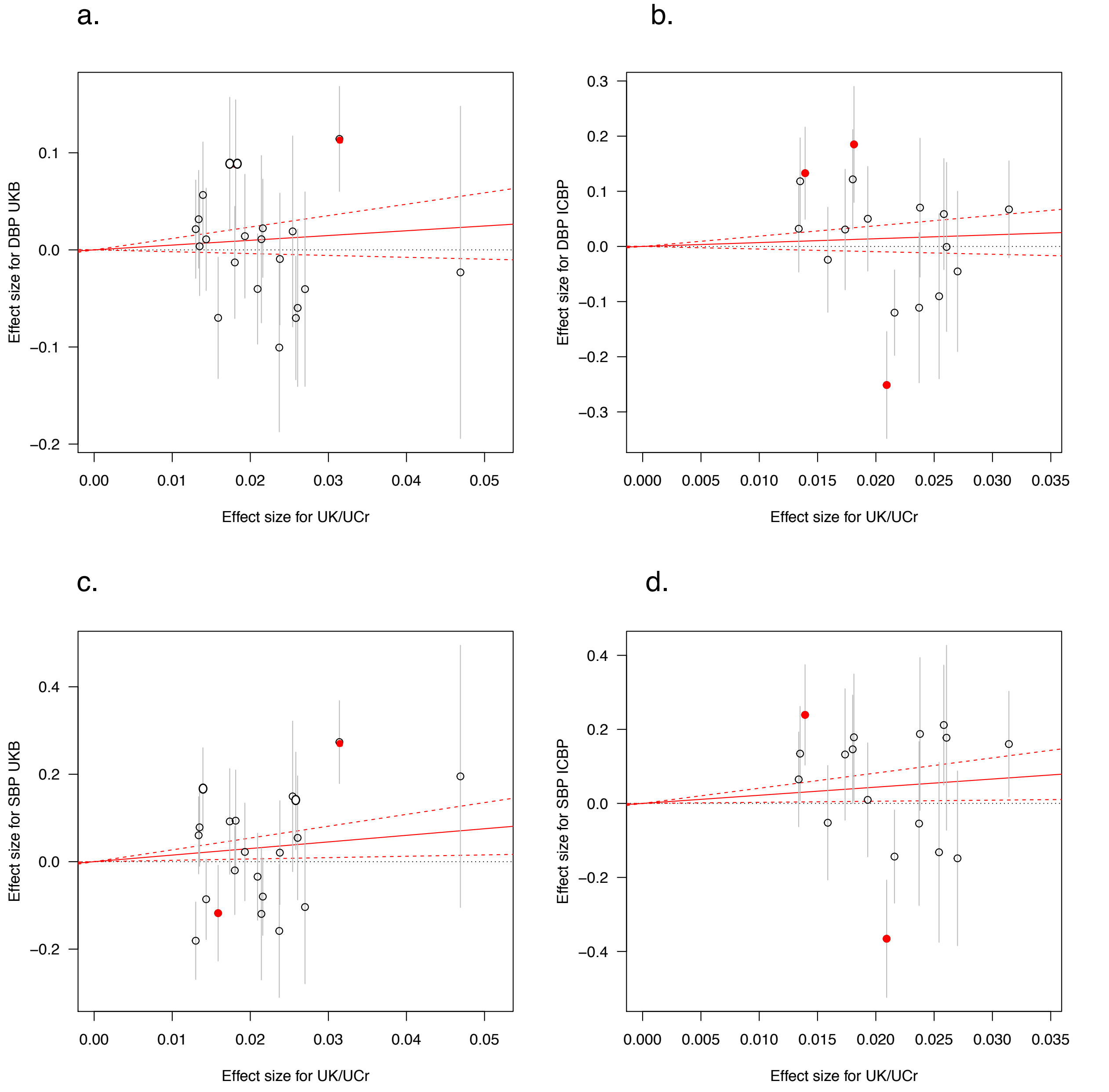

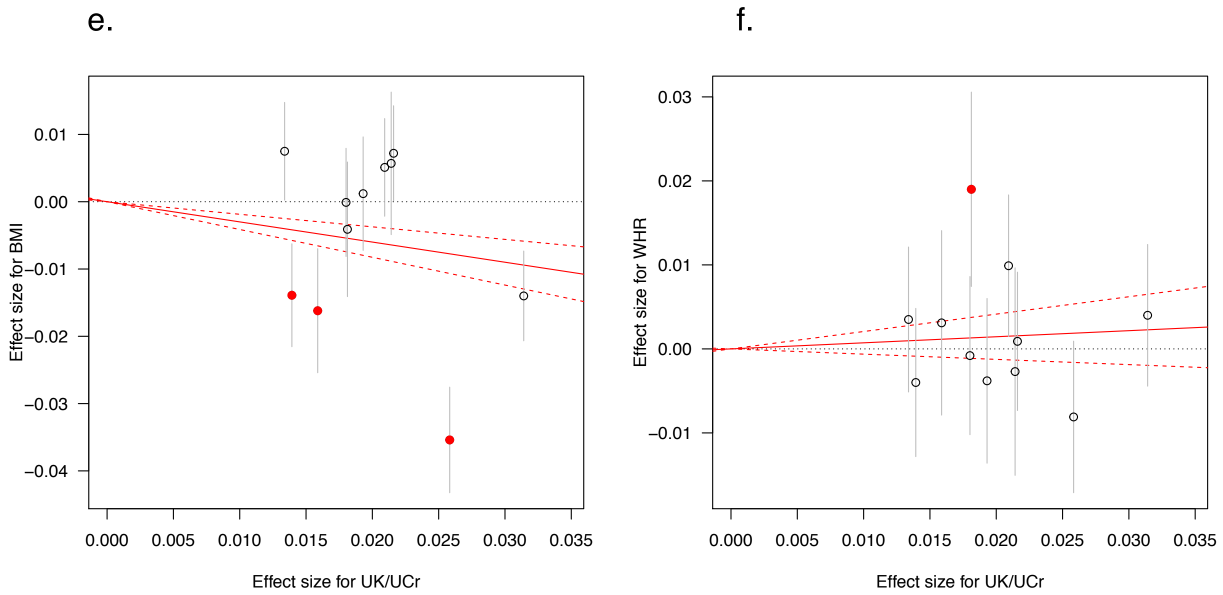
e.
f.
Effect size for BMI
0.01
0.00
−0.01
−0.02
−0.03
−0.04
0.000 0.005 0.010 0.015 0.020 0.025 0.030 0.035
Effect size for UK/UCr
Effect size for WHR
0.03
0.02
0.01
0.00
−0.01
0.000 0.005 0.010 0.015 0.020 0.025 0.030 0.035
Effect size for UK/UCr

**eFigure 8.** Effects of genetic variants on urinary albumin to creatinine ratio (UAlb/UCr) and risk of a. systolic blood pressure in UK Biobank (SBP UKB); b. systolic blood pressure from International Consortium of Blood Pressure (SBP ICBP); c. diastolic blood pressure in UK Biobank (DBP UKB); d. diastolic blood pressure from International Consortium of Blood Pressure (DBP ICBP); e. body mass index (BMI); f. waist-to-hip ratio (WHR); g. type 2 diabetes (T2D); h. atrial fibrillation (AF). Variants showing significant heterogeneity are highlighted in red.

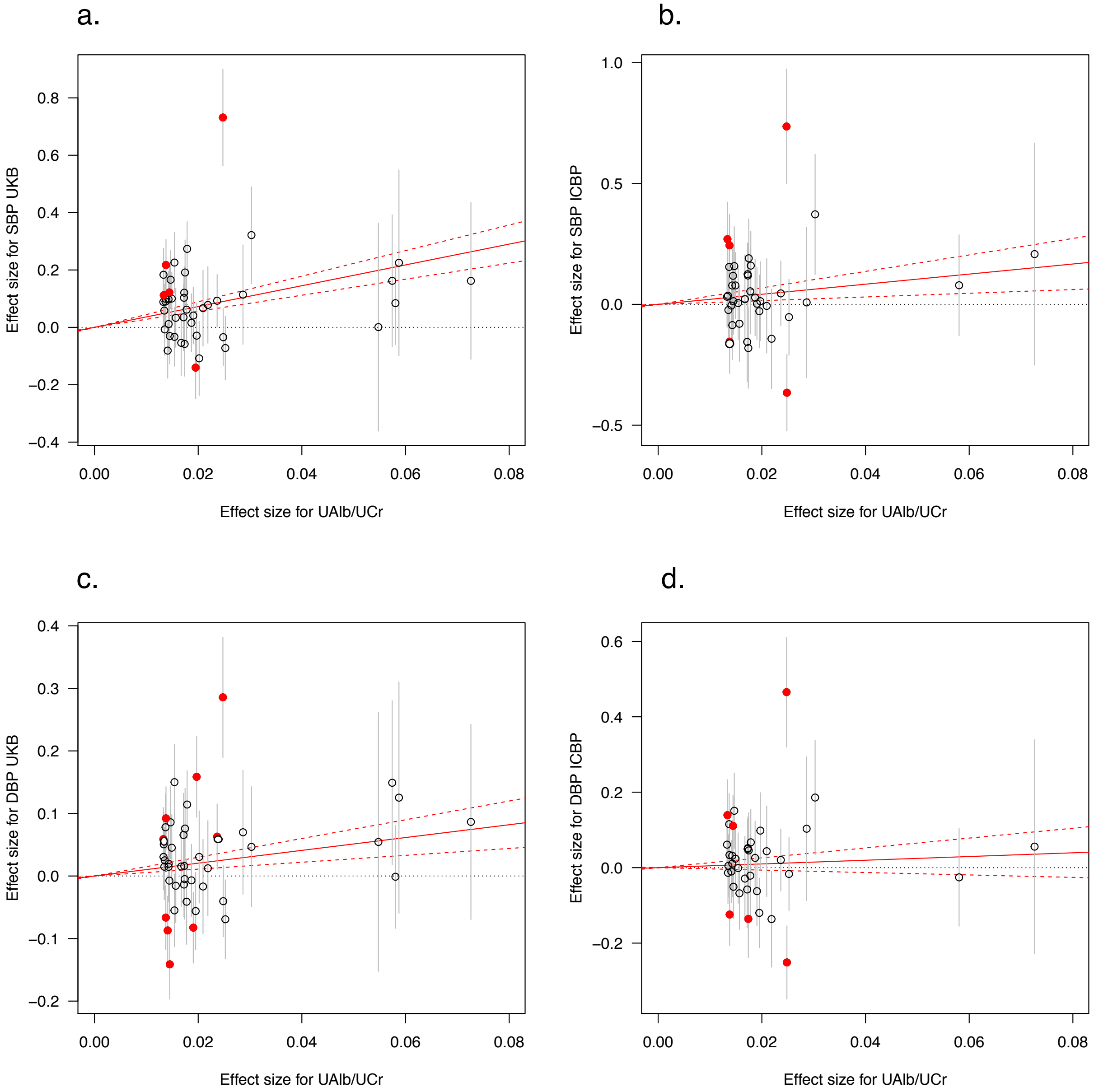

e.

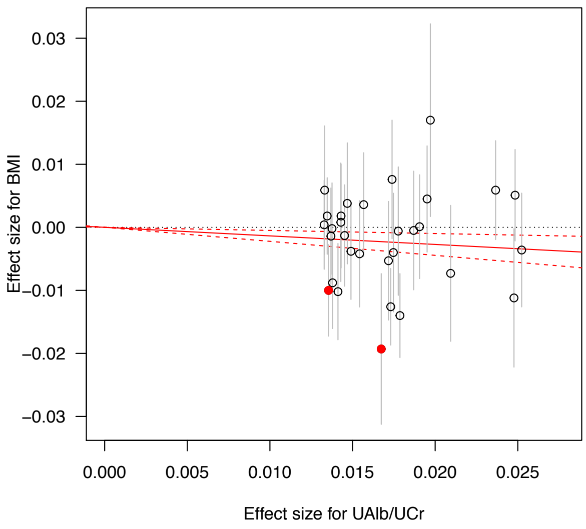
Effect size for BMI
0.03
0.02
0.01
0.00
−0.01
−0.02
−0.03
0.000
0.005
0.010
0.015
0.020
0.025
Effect size for UAlb/UCr

f.

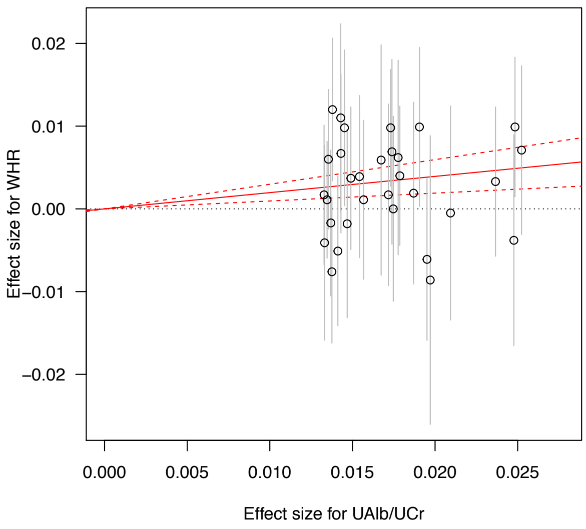
Effect size for WHR
0.02
0.01
0.00
−0.01
−0.02
0.000
0.005
0.010
0.015
0.020
0.025
Effect size for UAlb/UCr

g.

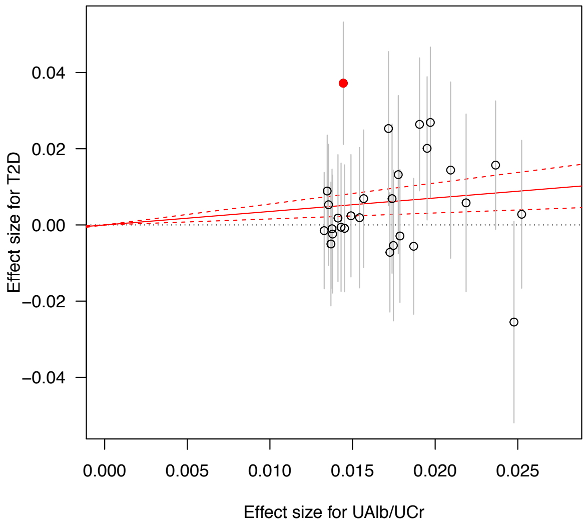
Effect size for T2D
0.04
0.02
0.00
−0.02
−0.04
0.000
0.005
0.010
0.015
0.020
0.025
Effect size for UAlb/UCr

h.

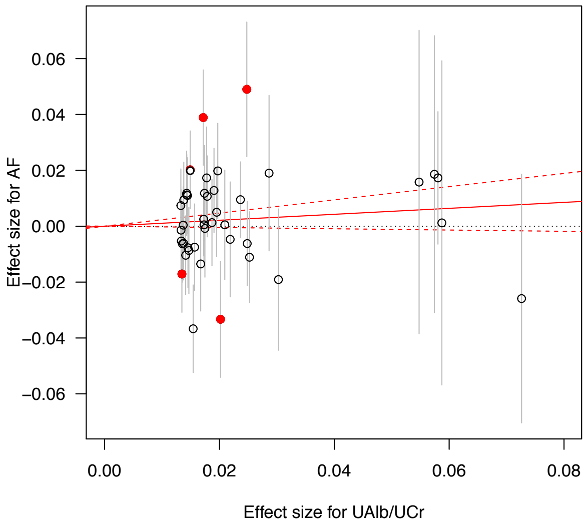
Effect size for AF
0.06
0.04
0.02
0.00
−0.02
−0.04
−0.06
0.00
0.02
0.04
0.06
0.08
Effect size for UAlb/UCr
